# TOM20 Gates PINK1 Activity And Mediates Its Tethering Of The TOM And TIM23 Translocases Upon Mitochondrial Stress

**DOI:** 10.1101/2023.08.07.552252

**Authors:** Mohamed A. Eldeeb, Andrew N. Bayne, Armaan Fallahi, Thomas Goiran, Emma J. MacDougall, Andrea Soumbasis, Cornelia E. Zorca, Jace-Jones Tabah, Rhalena A. Thomas, Nathan Karpilovsky, Meghna Mathur, Thomas M. Durcan, Jean-François Trempe, Edward A. Fon

## Abstract

Mutations in PTEN-induced putative kinase 1 (PINK1) cause autosomal recessive early onset Parkinson disease (PD). PINK1 is a Ser/Thr kinase that regulates mitochondrial quality control by triggering mitophagy mediated by the ubiquitin ligase Parkin. Upon mitochondrial damage, PINK1 accumulates on the outer mitochondrial membrane (OMM) forming a high molecular weight complex with the translocase of the outer membrane (TOM). PINK1 then phosphorylates ubiquitin, which enables recruitment and activation of Parkin followed by autophagic clearance of the damaged mitochondrion. Thus, Parkin-dependent mitophagy hinges on the stable accumulation of PINK1 on the TOM complex. Yet, the mechanism linking mitochondrial stressors to PINK1 accumulation and whether the translocases of the inner membrane (TIMs) are also involved, remain unclear. Herein, we demonstrate that mitochondrial stress induces the formation of a PINK1-TOM-TIM23 supercomplex in human cultured cell lines, dopamine neurons, and midbrain organoids. Moreover, we show that PINK1 is required to stably tether the TOM to TIM23 complexes in response to stress, such that the supercomplex fails to accumulate in cells lacking PINK1. This tethering is dependent on an interaction between the PINK1 NT-CTE module and the cytosolic domain of the Tom20 subunit of the TOM complex, the disruption of which, by either designer or PD-associated PINK1 mutations, inhibits downstream mitophagy. Together, the findings provide key insight into how PINK1 interfaces with the mitochondrial import machinery, with important implications for the mechanisms of mitochondrial quality control and PD pathogenesis.

## INTRODUCTION

Parkinson’s disease (PD) is a progressive neurodegenerative condition characterized by the selective loss of dopaminergic neurons in the substantia nigra of the midbrain (1). Importantly, current treatments only address the symptoms caused by the ensuing dopamine (DA) deficiency, but not the underlying molecular mechanisms that lead to neurodegeneration (1, 2). Analyses of the genes that cause inherited forms of PD point to mitochondrial dysfunction as a major contributor to the etiology of PD (2–4). An inherent challenge that mitochondria continuously encounter is the exposure to diverse stresses including high levels of reactive oxygen species and protein misfolding, which increase their likelihood of dysfunction (2, 5, 6). In response, eukaryotic cells have evolved an elaborate series of quality control mechanisms to identify, repair and/or eliminate defective mitochondria (7–11). One such mechanism is PINK1/Parkin mitophagy (selective degradation of mitochondria by autophagy), a process which involves PINK1, a mitochondrial Ser/Thr kinase (12) and Parkin, an E3 ubiquitin (Ub) ligase, encoded respectively by *PINK1* and *PRKN*, two genes in which loss-of-function mutations cause autosomal recessive early-onset PD (2, 12, 13). PINK1 acts upstream of Parkin and is essential for the mitochondrial localization and activation of Parkin. Upon mitochondrial damage, PINK1 builds up on the outer mitochondrial membrane (OMM) (14–21) where it phosphorylates both Ub and the Ub-like domain of Parkin (17, 22). Activated Parkin then ubiquitinates numerous OMM proteins, notably Mfn2 and VDAC (23–25), to initiate mitophagy (21, 22, 26). Thus, PINK1 acts a damage sensor on mitochondria, orchestrating the clearance of unhealthy mitochondria by Parkin (2, 21, 22, 27).

The PINK1 protein consists of an amino-terminal mitochondrial targeting sequence, a transmembrane helix and a 52 kDa cytosolic kinase domain. The cytosolic fragment alone carries Ub kinase activity and bears features of protein kinases such as catalytic and activation loops (21, 22, 28). PINK1 mRNA is co-transported with mitochondria in neurons, and the translated PINK1 protein is imported through the translocase of the outer membrane (TOM) complex, a large multimeric channel that is critical for the import of mitochondrial precursor proteins (2, 21, 29). Following import, PINK1 is cleaved at its N-terminus by MPP in the matrix and PARL in the inner mitochondrial membrane (IMM) (30, 31). While the mechanisms of PINK1 translocation to the PARL active site are still unknown, it has been proposed that AFG3L2, an IMM AAA+ protease, may play a role in the membrane dislocation of the PINK1 transmembrane domain (TMD) (a.a. 94-110) into a PARL-competent orientation (30, 32). Upon PARL cleavage between A103 and F104, PINK1 is retro-translocated to the cytosol where it is rapidly degraded by the proteasome-dependent N-degron degradation machinery (21, 32, 33). Thus, at steady-state, PINK1 levels are kept low, as corroborated by PINK1 half-life measurements around 30 minutes (34, 35). Remarkably, upon mitochondrial depolarization and import failure, commonly induced by treatment of cells with the protonophore cyanide m-chlorophenyl hydrazone (CCCP), PINK1 is no longer cleaved by proteases and builds up as a high-molecular weight (HMW) 720-800 kDa complex associated with the TOM machinery at the OMM (18, 19). The stabilization of PINK1 hinges on a motif of negatively charged residues (ie. E112, E113, E117) at the beginning of the N-terminal (NT) helix located between the TMD and the kinase domain, as well as the Tom7 subunit of the TOM complex (32, 36). Our group solved the crystal structure of the entire cytosolic domain of PINK1, revealing that the NT helix forms a module with its C-terminal extension (CTE), which is critical for PINK1 stabilization on TOM and subsequent activation by *trans* autophosphorylation on Ser228 (37). The NT-CTE module harbors PD mutations, which impair PINK1 activation and Parkin recruitment to mitochondria (32, 37, 38), thus highlighting the importance of characterizing this module.

The determinants for PINK1 import and accumulation at the IMM remain less defined: while most matrix-destined mitochondrial proteins rely on the translocase of the inner membrane (TIM23) complex and matrix import motor (primarily Tim44 and mtHsp70), there have been conflicting results on whether Tim23 knockdown affects PINK1 stabilization, PARL cleavage, and/or PINK1-TOM complex formation (32, 39). While it has been shown that TIM23-dependent substrates are typically transferred upon exiting the Tom40 pore to Tim50 (the primary receptor of the TIM23 complex which contains a presequence binding domain exposed to the IMS) (40–44), the potential interplay between PINK1 and Tim50 remains poorly characterized. It has also been reported that the IMM metalloprotease OMA1 can cleave PINK1 in depolarized or Tom7 knockout (KO) mitochondria to degrade misassembled PINK1 and prevent its aberrant hyperaccumulation (32, 39). Therefore, the mechanism of PINK1 association with the mitochondrial protein import machinery needs to be further elucidated.

Herein, we demonstrate that, in response to two distinct mitochondrial stressors, depolarization and misfolded proteins, PINK1 forms a supercomplex with the TOM and TIM23 complexes in multiple human cell types, including PD-relevant induced pluripotent stem cell (iPSC)-derived DA neurons and midbrain organoids. Using affinity purification-mass spectrometry in combination with AlphaFold, we identify an interaction between NT-CTE module of PINK1 and the Tom20 subunit of the TOM complex, which we show is required for PINK1-TOM-TIM23 supercomplex assembly, PINK1 kinase activation and downstream Parkin-mediated mitophagy. Importantly, PD-associated PINK1 mutations within this NT-CTE:Tom20 interface interfere with PINK1 supercomplex assembly and downstream mitophagy. Finally, we show that, in the absence of PINK1, TOM and TIM23 fail to assemble into a stable supercomplex. Thus, our work positions PINK1 as the first endogenous import substrate required to tether the TOM and TIM23 complexes into a stable supercomplex in response to mitochondrial stress in mammalian cells.

## RESULTS & DISCUSSION

### PINK1 assembles into a high molecular weight complex upon mitochondrial depolarization in human dopamine neurons and midbrain organoids

To validate previous reports of HMW PINK1 complex assembly upon depolarization and to investigate the dynamics of PINK1 accumulation, we transfected full-length PINK1 with either a C-terminal 3×FLAG tag or 6X-HIS fusion tag in HEK293T cells, treated with 20 µM CCCP or DMSO and monitored HMW PINK1 complex formation at various time points by Blue Native PAGE (BN-PAGE) (45). As predicted, both endogenous and the different tagged-PINK1 proteins assembled into a complex with an approximate molecular mass of 720 kDa **(Figure 1A)**. This complex likely contains the core TOM complex, as the endogenous Tom40 subunit can be observed to shifts its migration pattern upon CCCP treatment from 450 kDa, the approximate mass of the native TOM complex, to comigrate with PINK1 at 720 kDa **(Figure 1B)**. Proteinase K treatment led to the degradation of the HMW PINK1 complex, but not the inner membrane protein Tim22 **(Figure S1)**, indicating that the 720 kDa complex is exposed to the cytosolic side of the OMM in depolarized mitochondria. As reported previously (18), brief washout of CCCP to re-establish ΔΨm led to the disassembly of the endogenous HMW PINK1 complex, re-accumulation of the native 450 kDa TOM complex and degradation of PINK1 **(Figure 1B)**.

**Figure 1.**
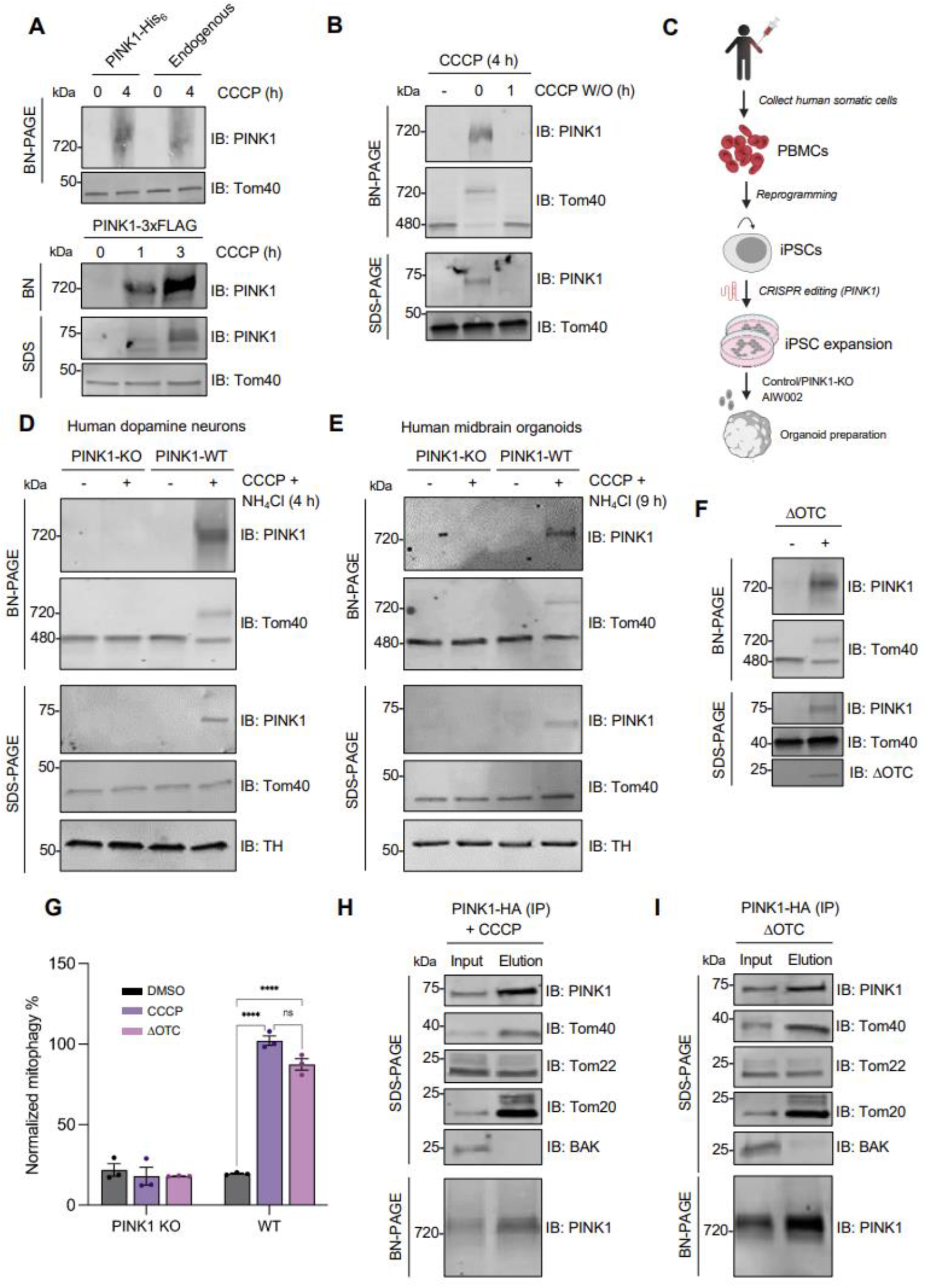
HMW PINK1 complex formation is ubiquitous across cell types and assembles in response to depolarization or misfolded protein accumulation. A) HEK293T cells (endogenous or expressing PINK1-His_6_ or PINK1-3xFLAG) were treated with 20 µM CCCP or DMSO for the indicated time points, lysed in 1 % digitonin solubilization buffer or SDS sample buffer, subjected to BN-PAGE or SDS-PAGE, and immunoblotted. B) U2OS cells expressing endogenous PINK1 were treated with 20 µM CCCP or for 4 h before washing out CCCP for the indicated times. Cells were lysed and immunoblotted for PINK1 and Tom40 as in Figure 1A. C) Schematic of human midbrain organoid generation from peripheral blood mononuclear cells (PBMCs) which have been reprogrammed into induced pluripotent stem cells (iPSCs) and CRISPR edited for PINK1 deletion. D) 6-week-old PINK1 KO dopamine neurons were treated with 20 µM CCCP or DMSO, subjected to BN-PAGE or SDS-PAGE, and then immunoblotted. E) 8.5-weeks Human midbrain organoids (PINK1-KO or WT) were treated with 20 µM CCCP and 5 mM ammonium chloride to inhibit lysosomal degradation as indicated, subjected to BN-PAGE and SDS-PAGE, and were immunoblotted using PINK1 and Tom40 antibodies. F) U2OS cells were transiently transfected with vector or ΔOTC constructs for 48 h and then fractionated. Mitochondrial-enriched fractions were solubilized, used for BN-PAGE (upper panels) and SDS-PAGE (lower panels), and immunoblotted. G) Quantification of mitophagy using the mt-Keima reporter assay in PINK1-HA transfected U2OS PINK1 KO cells following DMSO (black) or CCCP (violet) treatment, or ΔOTC transfection (pink). Bars indicate the relative level of mitophagy, normalized to WT PINK1 treated with CCCP, plotted as mean (n = 3) ± SEM. Two-way ANOVA with Tukey’s post hoc tests (n = 3), \**P* < 0.05; \*\**P* < 0.01; \*\*\**P* < 0.001; \*\*\*\**P* < 0.0001; ns, not significant. H) U2OS PINK1 KO cells were transfected PINK1-HA or mock vector and treated with 20 µM CCCP for 4 h. Mitochondria were isolated and HA immunocapture was performed. Bound proteins were eluted with HA peptide and fractions were subjected to SDS-PAGE or BN-PAGE immunoblotting. I) Mock or PINK1-HA transfected U2OS PINK1 KO cells were transfected with ΔOTC for 36 h followed by mitochondrial isolation and immunocapture. Bound proteins were eluted and subjected to SDS-PAGE or BN-PAGE immunoblotting.

To further characterize the endogenous HMW PINK1 complex in more physiological and PD-relevant cellular models, we used human iPSC-derived DA neurons and human midbrain organoids (hMBOs) **(Figure 1C)** (46). As a control, we used an isogenically matched PINK1 KO line, generated in the same iPSC background (47). Using immunofluorescence, we found that more than 75% of the neurons from both WT and PINK1 KO iPSCs were tyrosine hydroxylase-positive, indicating that the loss of PINK1 did not affect neuronal differentiation into DA neurons **(Figure S2)**. Both WT and PINK1 KO DA neurons and hMBOs were treated with either DMSO or CCCP and ammonium chloride to halt lysosomal degradation and increase PINK1 signal. The appearance of a CCCP/NH_4_Cl-dependent, PINK1- and Tom40-positive 720 kDa HMW complex could be observed on BN-PAGE in 6-week-old WT DA neurons **(Figure 1D)** and 8.5-week-old hMBOs **(Figure 1E)** but not in PINK1 KO DA neurons or hMBOs. Interestingly, the mobility of Tom40 itself didn’t shift in response to CCCP in the absence of PINK1. This suggests that despite the arrest of ΔΨm-dependent import of a myriad of proteins into mitochondria, no other endogenous TOM substrate, at least in neural human tissue, can substitute for the loss of PINK1 in the assembly of a stable complex with TOM. More broadly, these results in human DA neurons and hMBOs further re-enforce the robustness of the observed HMW PINK1 complex across models, including those with high relevance to PD.

### Misfolded mitochondrial protein accumulation elicits PINK1-TOM complex formation

Besides mitochondrial uncoupler-mediated import arrest, other stressors such as misfolded mitochondrial protein accumulation, have been reported to induce PINK1 stabilization on the OMM (48). Specifically, expression of a deletion mutant of ornithine transcarbamylase (ΔOTC), a protein that misfolds and accumulates in the mitochondrial matrix and induces the mitochondrial unfolded protein response (UPR^mt^) in mammalian cells, leads to PINK1 accumulation and Parkin recruitment on the OMM without disruption of membrane potential (48, 49). We hypothesized that ΔOTC expression would also result in the assembly of the HMW PINK1 complex. To address this, U2OS PINK1 KO cells expressing PINK1-HA were transiently transfected with a plasmid encoding ΔOTC, followed by mitochondrial extraction and BN-PAGE or SDS-PAGE. Similar to CCCP treatment, ΔOTC expression induced the formation of a 720 kDa HMW PINK1 complex **(Figure 1F)**. Moreover, both CCCP and ΔOTC elicited comparable levels of PINK1-dependent mitophagy **(Figure 1G),** as measured by flow-cytometry using the mt-Keima reporter (50). Using HA-immunopurification from mitochondrial lysates extracted from U2OS PINK1 KO cells expressing PINK1-HA, either treated with CCCP **(Figure 1H)** or co-expressing ΔOTC **(Figure 1I)**, we found that both stressors induced similar co-elution profiles and comparable enrichment patterns of TOM complex subunits (specifically Tom20, Tom40, Tom22). However, in contrast to CCCP, ΔOTC did not affect ΔΨm **(Figure S3)**, indicating that the two mitochondrial stressors triggered mitophagy via distinct mechanisms. Taken together, the findings demonstrate that ΔOTC induction of the UPR^mt^ results in a HMW PINK1 complex that contains a similar assortment of TOM subunits and appears competent to trigger mitophagy without requiring mitochondrial depolarization induced by CCCP.

### PINK1 forms a supercomplex with TOM and TIM23 upon mitochondrial depolarization

Previously, using a candidate approach, the HMW PINK1 complex was shown to contain components of the 450 kDa TOM complex (Tom40, Tom20, Tom22, Tom5, Tom6, Tom7 and Tom70) (18). Using an unbiased mass spectrometry-based approach, we sought to determine whether other components were part of the HMW PINK1 complex. Briefly, we isolated mitochondria from either mock or PINK1-2xStrep-His transfected, CCCP-treated HEK293T cells, affinity-purified PINK1, and subjected the elutions to LC-MS/MS analysis. Label-free quantification was performed to compare the enrichment in PINK1 versus mock-transfected cells **(Figure 2A, Table S1)**. First, our analysis re-affirmed the presence of previously known TOM subunits within the PINK1-TOM complex, as seen by the significant enrichment of Tom40, Tom22, Tom20, Tom5, Tom6, and Tom7 in the PINK1-transfected samples. Strikingly, three subunits of the TIM23 complex were also enriched within the CCCP-induced supercomplex, namely Tim50, Tim23, and Tim17B. This finding suggests that when PINK1 accumulates on mitochondria, it spans the entire mitochondrial intermembrane space and tethers the TOM and TIM23 complexes as part of a PINK1-TOM-TIM23 supercomplex. Indeed, upon CCCP treatment, both endogenous Tim50 and Tim23 co-migrated with PINK1 at 720 kDa on BN-PAGE consistent with the existence of a supercomplex containing PINK1 and components of TOM and TIM23 **(Figure 2B)**. In contrast, Tim22, a subunit of a distinct translocase complex (TIM22) at the IMM did not co-migrate with PINK1, TOM or TIM23 subunits, indicating that supercomplex does not include TIM22. Next, we overexpressed Tim50-FLAG or Tim23-FLAG and performed a FLAG immunopurification in CCCP-treated cells to further validate the components of the supercomplex. As expected from the LC-MS/MS data, we co-purified PINK1 along with Tom40 and Tom20 with both Tim50 **(Figure 2C)** and Tim23 **(Figure S4)**. Tim50 and Tim23 could reciprocally co-purify each other but not the Tim22 subunit of the IMM TIM22 complex. While Tom70 may play a role in steady-state PINK1 import (51, 52), our LC-MS/MS data indicate that Tom70 is not part of the PINK1-TOM-TIM23 supercomplex, consistent with a previous antibody-based gel shift study (19). Similar to our results with the TOM complex, none of the examined TIM23 subunits’ mobility shifted in response to CCCP in the absence of PINK1 (**Figure 2B**), again consistent with PINK1 functioning as an essential import substrate, required for the assembly of a HMW supercomplex in response to stress in mammalian cells.

**Figure 2.**
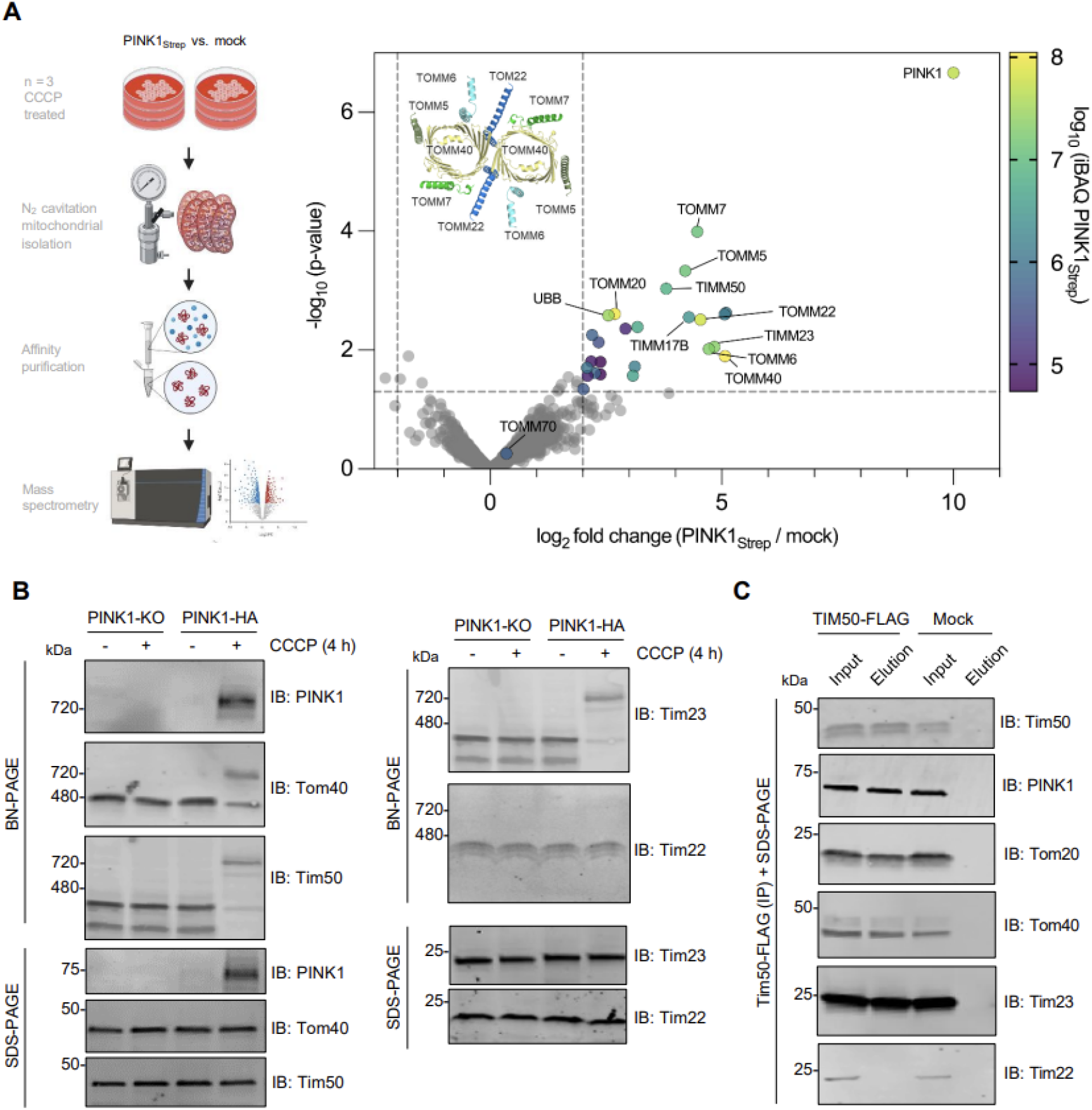
PINK1 forms a supercomplex with TOM and TIM23 complexes upon depolarization. **A)** HEK293T cells were transfected with PINK1-Strep or mock vector, mitochondria were isolated using nitrogen cavitation, StrepTactin purification was performed, and eluates were subjected to mass spectrometry analysis (left). Label-free quantification was performed (n = 3) and results are plotted as a volcano plot. Significantly enriched hits (-log_10_ p-value > 1.3 and log_2_ fold change > 2, indicated by dashed lines) are coloured according to their average iBAQ intensity across PINK1-Strep samples and selected hits are labelled. The core human TOM complex (PDB: 7CK6) is depicted as cartoons and labelled for reference. **B)** Mock or pCMV(d1) PINK1-HA transfected U2OS PINK1 KO cells were treated with 20 µM CCCP or DMSO as indicated, subjected to BN-PAGE or SDS-PAGE and immunoblotted as indicated. **C)** Mock or Tim50-FLAG transfected HEK293T cells were treated with 20 µM CCCP for 4 h followed by mitochondrial isolation and FLAG immunocapture. Bound proteins were eluted with FLAG peptide and fractions were subjected to SDS-PAGE immunoblotting.

### Tom20 directly interacts with PINK1 and gates downstream PINK1 activity

To provide structural context for our mass spectrometry data, we ran AlphaFold Multimer predictions iteratively using PINK1 as a bait against the top 30 of our significantly enriched mass spectrometry hits **(Figure S5)**. Strikingly, the highest scoring predictions from our unbiased screen were for complexes of PINK1:Ub and PINK1:Tom20. As the Ub binding site on insect PINK1 was previously determined (28), we decided to focus on the PINK1:Tom20 interaction, as it was the only TOM subunit previously shown to crosslink to PINK1 and form a direct interaction as part of the HMW PINK1 complex (18). Our AlphaFold prediction for PINK1 (95–581) and the cytosolic domain of Tom20 (51–145) also produced a high-confidence model, with all top 5 ranks converging on the same solution, as visualized by the low error scores in the predicted aligned error (PAE) plot **(Figure 3A)**. In this model, PINK1 binds to the Tom20 pre-sequence binding groove via its NT helix (a.a. 118-135) and αK helix (a.a. 524-544 in the CTE) **(Figure 3B)**. It is known that Tom20 is a labile component of the TOM complex with high lateral mobility, suggesting that it could dissociate and re-associate with the core TOM complex depending on the presence of presequences at the TOM gate (53). This model is also consistent with the recent cryo-EM structures of human TOM in which Tom20 co-purified with the TOM complex without chemical crosslinking, but only generated resolvable electron density upon crosslinking to Tom40 and Tom22 (54, 55). In our model, PINK1 accumulation could stabilize the interaction of Tom20 with the rest of the core TOM complex. Conversely, we hypothesized that the binding of PINK1 to Tom20 upon depolarization and the formation of a stable Tom20:PINK1 complex at the OMM would stabilize PINK1 folding on the OMM, effectively gating downstream PINK1 activity and mitophagy.

**Figure 3.**
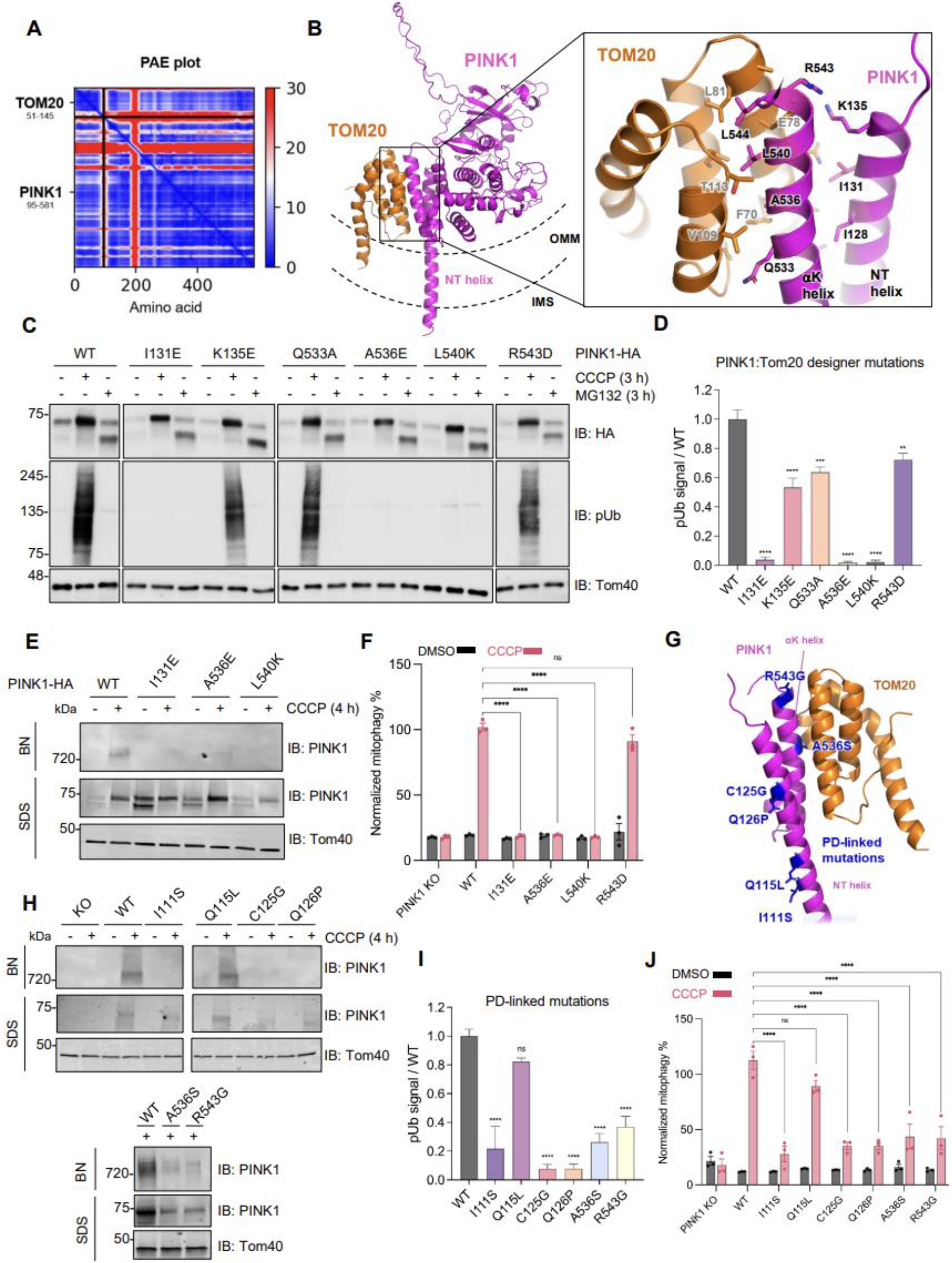
Characterization of the PINK1:Tom20 binding interface using designer and PD-linked mutations within the PINK1 NT-CTE. **A)** AlphaFold multimer v3 was run on the cytosolic domains of PINK1 (a.a. 95-581) and Tom20 (a.a. 51-145) with 20 recycles and an RMSD tolerance between cycles of 0.5 Å. The top ranked model was chosen for predicted aligned error plot (PAE) visualization. **B)** Structural visualization of the PINK1:Tom20 AlphaFold model. The top ranked model was AMBER relaxed and the binding interface was manually visualized and annotated using PyMOL. PINK1 and Tom20 are depicted in the context of the outer mitochondrial membrane (OMM) and intermembrane space (IMS) topology. **C)** U2OS PINK1 KO cells were transfected with pCMV(d1) PINK1-HA (WT or indicated mutants) and were treated with 20 μM CCCP or 10 µM MG132 for 4 h. Lysates were run on SDS-PAGE and immunoblotted as indicated. **D)** Quantification of immunoblots by densitometry for Fig. 3C. Band intensities were normalized relative to the level of Tom40 in the CCCP-treated sample, and the ratio to pSer65 of WT CCCP-treated sample was calculated. Bars represent mean ± SEM (n = 3). One-way ANOVA with Dunnett’s post-hoc test was performed (\**P* < 0.05; \*\**P* < 0.01; \*\*\**P* < 0.001; \*\*\*\**P* < 0.0001; ns, not significant). **E)** U2OS PINK1 KO cells were transfected with pCMV(d1) PINK1-HA WT or mutants and treated with 20 μM CCCP for 4 h. Mitochondria were extracted, solubilized, and analyzed via BN-PAGE or SDS-PAGE immunoblotting. **F)** U2OS cells transfected with pCMV(d1) PINK1 (WT or mutants) were treated with 20 μM CCCP or DMSO for 4 h and mitophagy was quantified using the mt-Keima reporter assays. Bars indicate the relative level of mitophagy, normalized to WT PINK1 treated with CCCP, plotted as mean (n = 3) ± SEM. Two-way ANOVA with Tukey’s post hoc tests (n = 3), \**P* < 0.05; \*\**P* < 0.01; \*\*\**P* < 0.001; \*\*\*\**P* < 0.0001; ns, not significant. **G)** PD-linked mutations visualized within the PINK1 NT-CTE and TOM20 AlphaFold model. **H)** U2OS PINK1 KO cells were transfected with pCMV(d1) PINK1-HA (WT or indicated mutants) and treated with 20 μM CCCP for 4 h. Mitochondria were extracted, solubilized, and analyzed by immunoblotting. **I)** U2OS PINK1 KO cells were transfected with pCMV(d1) PINK1-HA (WT or indicated mutants) and treated with 20 μM CCCP for 4 h. Lysates were immunoblotted and quantified as in Figure 3D. **J)** U2OS PINK1 KO cells were transfected with pCMV(d1) PINK1 mutants and assayed using the mt-Keima reporter assay and quantified as in Figure 3F.

To test this experimentally, we introduced a series of charged residues in place of key NT and αK residues that were predicted to interact with Tom20 in our AlphaFold model **(Figure 3B)**. We then transfected these mutants into U2OS PINK1 KO cells and quantified the levels of Phospho-S65 Ub (pUb) as a readout of PINK1 activity. While all our constructs still expressed PINK1 that was localized to mitochondria, as indicated by the formation of the PARL-cleaved 52 kDa PINK1 fragment upon MG132 treatment **(Figure 3C)**, three mutations (I131E, A536E, and L540K) were completely deficient in pUb generation despite the CCCP-dependent accumulation of full length PINK1 on SDS-PAGE **(Figure 3C and 3D)**. These three same mutants also failed to support the assembly of the PINK1-TOM-TIM23 supercomplex on BN-PAGE **(Figure 3E)** and were deficient in downstream mitophagy, as measured via the mt-Keima reporter assay **(Figure 3F)**.

In order to substantiate the relevance of this PINK1:Tom20 module in the context of PD, we also tested six PD-linked mutations (56) or naturally-occurring variants **(Figure 3G)** within the NT and αK helices for their effects on PINK1-TOM-TIM supercomplex assembly, PINK1 pUb generation, and downstream mitophagy. I111S, C125G, Q126P are located in the NT helix and are likely pathogenic for PD (57–59). Q115L is also in the NT helix, but is equally frequent in PD patients and controls, suggesting it is benign (60). Finally, the ClinVar database lists two variants of uncertain significance located in the CTE, A536S and R543G, which could impact binding to Tom20 (61). Five (I111S, C125G, Q126P, A536S, R543G) of the six mutants were deficient in forming the PINK1-TOM-TIM23 supercomplex **(Figure 3H)**. Moreover, these same five PD mutants were deficient in pUb generation **(Figure 3I)** and in downstream mitophagy **(Figure 3J)**. We confirmed that I111S, C125G and Q126P exhibited a severe defect in Parkin recruitment to mitochondria (**Figure S6**) and a reduced colocalization with mitochondria on confocal immunofluorescence microscopy (**Figure S7**). Conversely, the benign variant Q115L could support Ub phosphorylation, PINK1-TOM-TIM23 supercomplex assembly and mitophagy, and only showed a slight delay in Parkin recruitment (**Figure S6**). Based on the structural model of PINK1-Tom20 **(Figure 3G)**, neither I111S nor Q115L would affect Tom20 binding, but I111S is located further upstream and could disrupt an interaction with other TOM subunits. Taken together, these findings underscore Tom20 as a crucial interactor of PINK1, acting as a ratchet that helps stabilize PINK1 on the TOM complex when import via TIM23 is stalled long enough to allow folding of the kinase domain via the NT-CTE module. This folding and stabilization of PINK1, in turn, would enable autophosphorylation, ubiquitin phosphorylation, and downstream mitophagy. These findings are consistent with previous results regarding the regulation of mitophagy through celastrol-mediated modulation of PINK1-TOM20 binding (62).

### PINK1 is required to tether the TOM and TIM23 complexes

The translocation of protein precursors into mitochondria requires the transfer of precursors from the TOM translocase at the OMM to the translocation machineries at the IMM, including the TIM23 complex, the main IMM translocase for presequence-containing precursors (63). Despite their localization on different mitochondrial membranes, under certain circumstances, usually in the context of overexpression of artificial or synthetic stalled import substrates in yeast, TOM and TIM23 have been shown to assemble into a supercomplex, tethered by the artificial substrate trapped in their import channels (64–66). Given our observation that the assembly of TOM and TIM23 into a stable supercomplex in response to mitochondrial stress requires PINK1 **(Figure 1D, 1E and 2B**), we asked whether PINK1 could serve as a *bone fide* native tethering import substrate in mammalian cells. To address this, we performed Tom22-FLAG and Tim50-FLAG immunopurifications under DMSO and CCCP conditions in both WT and PINK1 KO HeLa cells. As expected, Tom22 co-purified with Tom20, another subunit of the TOM complex, both at baseline and after CCCP treatment (**Figure 4A)**. In contrast, Tom22 only co-purified PINK1 and the TIM23 subunits, Tim23 and Tim50, upon depolarization with CCCP, confirming the requirement of mitochondrial stress for PINK1-TOM-TIM23 supercomplex assembly (**Figure 4A)**. Reciprocally, Tim50 co-purified the TIM23 subunit, Tim23, at baseline but only co-purified PINK1 and the TOM subunits, Tom 20 and Tom22, upon depolarization (**Figure 4B)**. Under none of the conditions did Tom22 or Tim50 co-purify the Tim22 subunit of the TIM22 translocase at IMM. Most importantly, in the absence of PINK1, neither Tom22 nor Tim50 co-purified subunits from the opposing complex, strongly arguing that PINK1 is required to tether the TOM and TIM23 complexes in response to mitochondrial stress **(Figure 4A and B)**. Next, we determined whether the Tom20:PINK1 NT-CTE interaction described above **(Figure 3)** was critical for PINK1 to tether the TOM and TIM23 complexes by performing PINK1-HA immunopurifications in CCCP-treated U2OS PINK1 KO cells transfected with either WT or the I131E, A536E, L540K, and R543D PINK1 mutants. As expected, Tom20 co-purification was abrogated in the three mutants (I131E, A536E, L540K) which showed deficient pUb generation, PINK1-TOM-TIM23 supercomplex formation and mitophagy **(Figure 3C-F)**, but not in R543D, a mutant whose function was similar to WT PINK1 **(Figure 4C)**. Furthermore, the defective PINK1 mutants also prevented the co-purification of Tom40, suggesting that the Tom20:PINK1 NT-CTE interaction is required for PINK1 to stably bind the TOM complex. Finally, even if these same three mutants were properly localized to mitochondria **(Figure 3C)**, they failed the co-purify the Tim50 subunit of the TIM23 complex, suggesting that Tom20:PINK1 NT-CTE binding is required for PINK1 to tether TOM and TIM23 into a supercomplex. We also performed a pulldown of these Tom20-binding deficient PINK1 mutants using Tim50-FLAG immunopurification (**Figure S8**). In the presence of these PINK1 mutants, Tim50-FLAG co-purified with Tim23 but not Tom40 or PINK1, confirming that stable TOM-TIM23 supercomplex formation hinges on the PINK1:Tom20 interaction. Next, we attempted to characterize how modulation of TIM23 complex levels would impact the PINK1-dependent tethering of TOM-TIM23. To this end, we knocked down Tim50 using CRISPRi or overexpressed Tim50-FLAG in CCCP-treated cells and visualized PINK1-TOM-TIM23 formation by BN-PAGE. Whereas Tim50 knockdown did not reduce PINK1 levels overall, as seen on SDS-PAGE, it did reduce the amount of PINK1-TOM-TIM23 supercomplex on BN-PAGE **(Figure 4D)**. Furthermore, overexpressing Tim50-FLAG leads to an apparent increase in PINK1-TOM-TIM23 supercomplex levels. It is critical to note that Tim50 knockdown also reduced the levels of the Tim23 subunit, which makes it difficult to draw conclusions on whether this reduction is Tim50 specific or simply due to disruption of the stoichiometry of the TIM23 complex. Still, these findings suggest that the entire TIM23 complex is one of the key factors for the stabilization of PINK1 on the TOM complex. In line with this, another recent study revealed that reduction of Tim23 levels also attenuated PINK1-TOM-TIM23 complex formation, though the effects of Tim50 knockdown on supercomplex assembly were not characterized (39). It was also proposed that PINK1 which escaped Tim23 binding was susceptible to OMA1 proteolysis, which suggests that the abundance of the TIM23 complex and its ability to bind PINK1 are critical for maintaining the stability of accumulated PINK1 within the supercomplex. Other studies have shown conflicting results that Tim23 and Tim50 knockdown did not affect overall PINK1 levels or its accumulation on SDS-PAGE following depolarization, though the consequences on supercomplex formation were not investigated (32). Future studies will be essential to clarify the specific roles of Tim23 and Tim50 in PINK1 accumulation and PINK1-TOM-TIM23 assembly.

**Figure 4.**
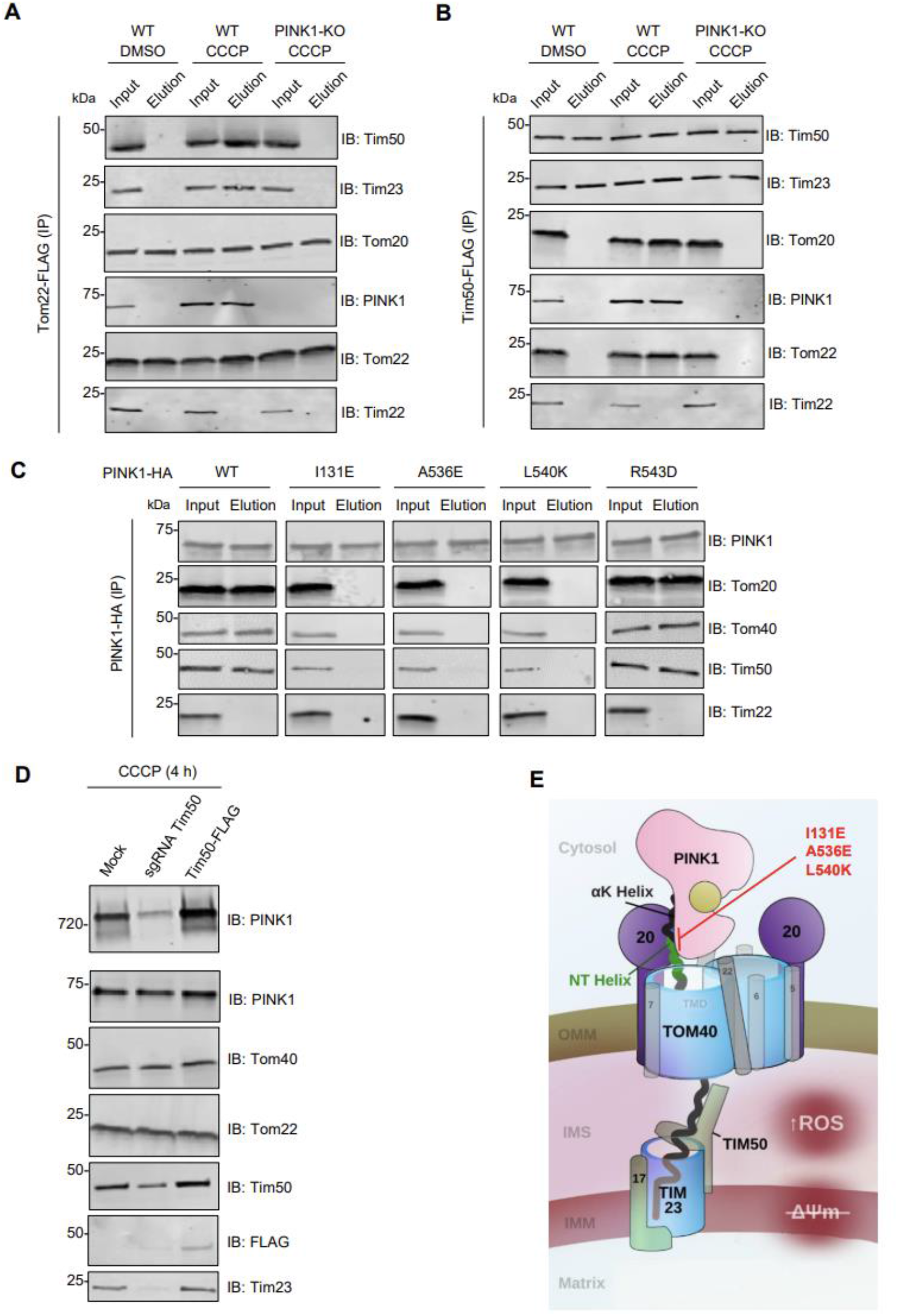
PINK1 endogenously tethers TOM and TIM23 following import arrest in a Tom20-dependent manner. **A)** U2OS cells (WT or PINK1-KO) were transfected with Tom22-FLAG and treated with 20 µM CCCP or DMSO for 4 h, followed by mitochondrial isolation and FLAG immunocapture. Bound proteins were eluted with FLAG peptides and subjected to SDS-PAGE immunoblotting. **B)** U2OS cells (WT or PINK1-KO) were transfected with Tim50-FLAG and treated with 20 µM CCCP or DMSO for 4 h, followed by mitochondrial isolation and FLAG immunocapture as in Figure 3A. **C)** U2OS PINK1 KO cells transfected with pCMV(d1) PINK1-HA (WT or Tom20 binding deficient mutants) were treated with 20 µM CCCP for 4 h followed by mitochondrial isolation and HA immunocapture. Bound proteins were eluted with HA peptides and subjected to SDS-PAGE immunoblotting. **D)** U2OS PINK1 KO cells were transfected with mock vector, CRISPRi sgRNA targeting Tim50, or Tim50-Flag, and treated with 20 µM CCCP or DMSO for 4 h. Lysates were subjected to SDS-PAGE or BN-PAGE immunoblotting as previously described. **E)** Simplified schematic of PINK1 topology within the PINK1-TOM-TIM supercomplex. Accessory TOM subunits are depicted as gray. PINK1 NT and αK helices are labelled along with Tom20 (depicted as “20”). This model of Tom20 binding places the transmembrane domain (TMD) of PINK1 in line with Tom40 in the OMM. For simplicity, the PINK1 N-terminus is depicted as passing through the Tom40 pore.

### Concluding remarks

Taken together, our findings establish that, in response to mitochondrial stress, PINK1 accumulates and tethers core components of the TOM and TIM23 complexes, leading to the assembly of a 720 kDa PINK1-TOM-TIM23 supercomplex **(Figure 4E)**. The complex forms across physiologically relevant cell models, including DA neurons and hMBOs and different stressors, including depolarization and the ΔOTC-induced UPR^mt^. These findings reinforce the broad significance of this biochemical assembly in neuronal mitochondria and the conjecture that the PINK1-TOM-TIM23 complex serves as an integrator of multiple mitochondrial stress signals. It will be crucial to further characterize the potential differences in subunit composition between different cell-types and between CCCP- and ΔOTC-induced PINK1 assemblies at the level of TIMs and the matrix import motor.

While TOM-TIM23 supercomplexes have been visualized previously in yeast using electron cryotomography (67) or crosslinking to a synthetic import intermediate (Jac1 fused to superfolding GFP) (64), PINK1 represents, to the best of our knowledge, the first endogenous physiological import substrate to tether and stabilize TOM-TIM23 supercomplex assembly with high occupancy following mitochondrial stress. Although the precise oligomeric states of the TOM and TIM23 machinery in complex with PINK1 remain to be determined, based on the apparent 720 kDa molecular mass for PINK1-TOM-TIM23 on BN-PAGE and the list of co-interactors from our mass spectrometry data, it is likely that the TOM machinery is present as a tetramer in its PINK1-bound state, as TOM dimers or trimers containing PINK1 and TIM23/17/50 heterotrimers should migrate at < 550 kDa in molecular mass. This would be consistent with the observation of TOM tetramers by cryoelectron microscopy, in both yeast and human purified TOM complexes (55, 68). To this end, deciphering the precise stoichiometry of the HMW PINK1 complex will be crucial to understand the physiological ramifications of these mitochondrial assemblies.

TOM and TIM23 tethering by PINK1 clearly depends on the Tom20:PINK1 NT-CTE interaction on the OMM side of the supercomplex and possibly on Tim50 on the IMS/IMM side. While Tom20 is known to bind presequences to facilitate protein import, our work positions PINK1 as a non-canonical import substrate that bridges TOM and TIM23 complexes upon mitochondrial stress in a Tom20-dependent manner. Thus, we envision that in healthy mitochondria, competent for import, PINK1 would be efficiently pulled through the TOM complex via AFG3L2 or other TIM machinery before the NT-CTE had sufficient time to fold completely, effectively dislodging the Tom20 interaction to expose PINK1 to the PARL active site in the IMM for subsequent PINK1 cleavage and degradation. In contrast, in mitochondria in which import is compromised, the fully folded PINK1 NT-CTE would remain bound to Tom20 in an orientation where the PINK1 TMD is not accessible for cleavage, effectively allowing PINK1 to remain Tom20-bound, its kinase domain to fold properly at the cytosolic side of the OMM, and trigger downstream mitophagy. This model sets the stage for further studies regarding the mechanisms by which the mitochondrial import machinery sense misfolded protein-induced mitochondrial damage signals. Future studies will also be warranted to study the functional impact of PINK1 PD-mutations on the PINK1 complex formation in DA neurons and its subsequent impact on neuronal mitochondrial biology, a process which will likely provide key functional insights into the molecular basis of PD.

Overall, our work yields key insights into the determinants for PINK1-TOM-TIM23 supercomplex assembly and downstream mitophagy as well as shedding light on yet another aspect of how mitochondrial translocase’s fine-tune PINK1-mediated mitophagy, laying the foundation for future studies on mitochondrial quality control in PD biology.

## MATERIALS AND METHODS

### Cell culture and immunoblotting

U2OS PINK1 KO cell lines were generated by CRISPR as previously reported (37). Cells were cultured in DMEM (Dulbeco’s Modified Eagle Medium) supplemented with 10% fetal bovine serum and 1x Pen/Strep and grown at 37°C and 5% CO_2_. pCMV(d1)TNT PINK1(WT)-3HA, denoted as PINK1-HA throughout the manuscript, was obtained from Noriyuki Matsuda for attenuated PINK1 expression (69). PCR mutagenesis was used to generate mutations in the NT-CTE of human PINK1 in the pCMV(d1)TNT PINK1 vector. Other plasmids used include pReceiver M14 PINK1-3xFLAG (Genecopoeia), denoted as PINK1-3xFLAG, and pTT5-PINK1-2xStrep-His, denoted as PINK1-Strep or PINK1-HIS. U2OS PINK1 KO cells were transfected with 1 µg pCMV(d1)TNT PINK1(WT)-3HA or indicated plasmids in 6-well plates for 48 h, followed by treatment with 20 µM of CCCP or equal volume of DMSO for 4 h. Whole cells were harvested, resolved by SDS-PAGE and immunoblotted on nitrocellulose as described previously (70). The following antibodies were used: PINK1 D8G3 (Cell Signaling cat# 6946), Anti-Tom40 Antibody (H-7): sc-365466; and phospho-S65 Ub (Millipore EMD, cat# ABS1513), Anti-Tom70 Antibody (A-8): sc-390545; Anti-Tom22 Antibody (Sigma, cat# T6319); Anti-Tom20 antibody (Santa Cruz Biotechnology, sc-17764), Anti-HA 6E2 (Cell Signalling cat# 2367), Anti-Tim23 (abcam cat# ab230253), Anti-Tim22 (abcam, cat# ab251909) and Anti-Tim50 (santa Cruz Biotechnology, sc-393678), and Anti-BAK antibody (Cell Signaling, cat #3814). The membranes were blocked with 3% fish skin gelatin (Sigma) in 1X PBS with 0.1% Triton X-100), probed with primary and secondary antibodies, and imaged with an Odyssey infrared imaging system using the manufacturer’s recommended procedures (LI-COR). For Figure 3C and all phospho-S65 Ub (pUb) blots, proteins were resolved on 4-20% (for pUb) or 10% SDS-PAGE (for TOM40 and HA) gels, transferred for 90 mins at 250 mA onto polyvinylidene difluoride (PVDF) membranes, blocked in 1X TBS-T (0.1% Tween 20) with 5% BSA, and incubated with primary antibodies overnight at 4°C. The following day, membranes were washed, incubated with HRP-conjugated secondary antibody for 1 h at RT, washed again, and then visualized using ClarityTM chemiluminescence (Bio-Rad).

### Mitochondrial isolation and Blue-Native-Poly-Acrylamide Gel Electrophoresis (BN-PAGE)

U2OS PINK1 KO cells were transfected with pCMV(d1)TNT PINK1(WT)-3HA or the indicated mutants for 36 h, followed by treatment with 20 µM of CCCP or equal volume of DMSO for 4 h. The cells were then harvested in mitochondrial isolation buffer containing 20 mM HEPES (pH 7.4), 220 mM mannitol, 70 mM sucrose and cOmplete protease inhibitor cocktail EDTA-free (Roche). Nitrogen cavitation was performed to pellet mitochondria as described previously (37, 71). The mitochondria were then solubilized in a buffer containing 20 mM BIS-TRIS (pH 7.3), 100 mM NaCl, 10% glycerol, protease inhibitors and 1% digitonin and left on a rotor for 3 h at 4°C. The suspension was spun at 5000 g for 5 mins at 4°C and the supernatants were collected. BN-PAGE was performed using Native^TM^ PAGE Running Buffer (Invitrogen) containing 0.002% G-250 (Invitrogen). Gels were shaken in denaturation buffer (10 mm Tris-HCl pH 6.8, 0.1% SDS, and 0.006% 2-mercaptoethanol) for 60 mins after electrophoresis and then transferred to PVDF membranes for immunoblotting. The membranes were blocked with 3% fish skin gelatin (Sigma) in 1X PBS with 0.1% Triton X-100, probed with primary and secondary antibodies, and imaged with an Odyssey infrared imaging system (LI-COR).

### Mitochondrial GFP-Parkin recruitment time-lapse microscopy

U2OS PINK1 KO cells were seeded on a 35-mm Glass Bottom 4 Compartment Dish (Greiner Bio-One). Cells were co-transfected with WT eGFP-Parkin and pCMV(d1)TNT PINK1(WT)-3HA or mutants. 48 h after transfection, cells were stained with MitoTracker DeepRed FM (ThermoFisher) at a final concentration of 50 nM following manufacturer’s guidelines. Cells were then transferred to a heated stage maintained at 37 °C and 5% CO_2_ using a Zeiss temperature controller and cell perfusion system. Cells were treated with CCCP at a final concentration of 15 uM. Microscopy was performed on a Zeiss AxioObserver.Z1 inverted fluorescent microscope. Fully automated multidimensional acquisition was controlled using the Zen Pro software (Zeiss). Fixed exposure times were as follows: GFP 150 ms; and MitoTracker 40 ms. Images were taken at 4 min intervals for a total of 120 min. Parkin recruitment was visualized by the appearance of punctate GFP fluorescence overlapping Mitotracker fluorescence. The percentage of cells exhibiting Parkin recruitment to the mitochondria was calculated at 4 min intervals over 120 mins. Approximately 200 cells were examined per mutant over four separate and biologically independent experiments. The fluorescence intensity of GFP-positive puncta was not considered in Parkin recruitment analysis. For statistical analysis, a two-way analysis of variance (ANOVA) with Bonferroni post-test was performed, *P<0.05.

### Human midbrain organoid generation, mitochondrial extraction and blue-native PAGE

Human midbrain organoids (hMBOs) were derived from the healthy control cell line AIW002-02 reprogrammed from human peripheral blood monocytes into iPSCs (72) and an isogenic CRISPR-induced deletion of PINK1 in the AIW002-02 line (47). hHMBOs were generated from iPSC cells using Enuvio embryoid-body disks (EB-DIST) on day 0 and transferred to bioreactors after 8 days following the previously reported protocol (73). After 8.5-weeks, organoids were transferred to media containing 20 µM CCCP and 5 mM NH_4_Cl or equivalent DMSO for 9 h. After treatment, organoids were washed multiple times with PBS and then harvested in mitochondrial isolation buffer containing 20 mM HEPES (pH 7.4), 220 mM mannitol, 70 mM sucrose and cOmplete protease inhibitor cocktail EDTA-free (Roche). Nitrogen cavitation was performed on ice for 5 minutes at a pressure of 500 psi. The resulting suspension was centrifuged for 5 minutes at 3000 rpm and the pellet was resuspended in mitochondrial isolation buffer. Mitochondrial fractions were pelleted by centrifugation for at 12000 rpm for 20 minutes. The mitochondria were then solubilized in a buffer containing 20 mM BIS-TRIS (pH 7.3), 100 mM NaCl, 10% glycerol, protease inhibitors and 1% digitonin and left on a rotor for 3 hours at 4°C. The suspension was spun down at 5000 g for 5 minutes at 4°C and the supernatants were carried forward for BN-PAGE gels.

### Dopaminergic neuron generation from induced pluripotent stem cells

The previously validated AIW002 control and PINK1-KO pluripotent stem cell (iPSC) lines were used (47). IPSC culture and neuronal differentiation were conducted according to previously established protocols (74–76). Briefly, iPSCs were passaged using Relsr, plated on Matrigel coated dishes and maintained in mTeSR basal media. Midbrain neuronal precursor cells (NPCs) were generated following a previously established protocol. Briefly, iPSCs were grown until 70% confluent and then dissociated using gentle cell dissociation reagent and transferred to uncoated flasks in NPC Induction Media (Composed of DMEM/F12 supplemented with 1x N2, 1x B-27, 1x MEM NEAA solution, 200 ng/mL noggin, 200 ng/mL SHH, 3 μM CHIR-99021, 10 μM SB431542 and 100 ng/mL FGF-8) containing 10 μM of Y-27632 (ROCK inhibitor) to allow for embryoid body (EB) formation. Media was refreshed every 48 h and on the 7^th^ day, EBs were transferred to polyornithine/laminin coated flask, allowing EBs to attach to the culture surface. Media was again changed every 48 h and on the 7^th^ day EBs were dissociated into small colonies by trituration in gentle cell dissociation media, and replated on polyornithine/laminin coated flasks, splitting 1 flask of EBs into 2-3 flasks for expansion. The resulting NPC monolayer was grown again for 7 days in NPC induction media until 100% confluent, at which time NPCs were harvested with gentle cell dissociation media and frozen in FBS with 10 % DMSO.

For differentiation into dopaminergic neurons, NPCs were thawed in NPC maintenance media (composed of DMEM/F12 supplemented with 1x N2, 1x B-27, 1x MEM NEAA solution, 100 ng/mL FGF-8 and 2 μM purmorphamine), plated on polyornithine/laminin coated flasks and expanded for at least 7 days with media change every 24-48 h as needed. To plate NPCs for final differentiation into dopaminergic neurons, NPCs were dissociated using Accutase, counted and plated on polyornithine/laminin coated dishes at the desired density in dopaminergic differentiation media (Composed of Neurobasal A media supplemented with 1x B27, 1x N2, 1x Antibiotic-Antimycotic, 20 ng/mL BDNF, 20 ng/mL GDNF, 200 μM Ascorbic acid, 0.5 mM db-cAMP and 0.1 μM Compound E). After 5 days, media was supplemented with 0.1 μg/mL mitomycin C to remove any remaining proliferative cells. Dopaminergic neurons were maintained and matured by refreshing 1/3^rd^ of the culture media every 5-7 days.

For imaging experiments neurons were plated on 96-well plates at a density of 15,000 cells per well. For biochemical experiments neurons were plated on either 6-well plates at a density of 750,000 cells per well or 15 cm dishes at a density of 10 million cells per plate, and then were treated with 20 µM CCCP and 5 mM NH_4_Cl or DMSO for 4 h, harvested, lysed, and subjected to SDS-PAGE or BN-PAGE analysis in the same manner as U2OS PINK1 KO cells described above.

### Characterization of neurons by immunofluorescence

Neuronal identity was assessed by immunofluorescent staining for Map2 (neurons) and tyrosine hydroxylase (TH - dopaminergic neurons). After 4 weeks of differentiation neurons were fixed with 4% paraformaldehyde for 20 mins, permeabilized for 10 mins with 0.3% saponin in PBS, and blocked for 1 h with 1% BSA and 4% goat-serum in PBS. Primary antibodies against TH (Pelfreeze, P40101) and Map2 (Encor, CPCA-MAP2) were diluted (1:500 and 1:2000 respectively) in blocking solution and incubated at 4 degrees overnight. Neurons were then washed 3 times in PBS and incubated for 1 hour with AlexaFluor secondary antibodies in blocking solution. Neurons were then washed twice and stained with Hoechst before imaging.

Imaging was performed on an Opera Phenix high content confocal microscope using 20X water immersion objective. Image analysis was performed using the Columbus software and data processing was then conducted using R studio. Briefly, nuclei were first identified by the Hoechst channel, and surrounding somal area was identified by Map2 staining. TH labelling intensity was then quantified within this Map2-defined region. Single-cell data were then exported as text files and imported into R studio for processing. A pre-processing script was used to filter objects based on nuclear size, nuclear shape and Map2 staining intensity to identify neuronal cells, and TH positivity was used to define the percentage of dopaminergic neurons.

### Mitophagy assay using the mt-Keima reporter

Mitophagy was examined using a FACS-based analysis of mitochondrially targeted Keima (mt-Keima) as previously described in (50, 77, 78). Briefly, U2OS PINK1 KO cells stably expressing GFP-Parkin (WT) and ecdysone-inducible mt-Keima were transfected with PINK1-HA (WT or indicated mutants) for 36 h, induced with 10 μM ponasterone A, and treated with 20 μM CCCP for 4 h. For FACS analysis, cells were trypsinized, washed and resuspended in PBS prior to their analysis on a LSR Fortessa (BD Bioscience) equipped with 405 and 561 nm lasers and 610/20 filters. Measurement of lysosomal mitochondrially targeted mt-Keima was made using a dual excitation ratiometric pH measurement where pH 7 was detected through the excitation at 405 nm and pH 4 at 561 nm. For each sample, 10,000 events were collected and single, GFP-Parkin-positive cells were subsequently gated for mt-Keima. Data were analysed using FlowJo v10.1 (Tree Star). For statistical analysis, the data represent the average percentage of mitophagy from at least three independent experiments, and P values were determined by one-way ANOVA with Dunnett’s post-hoc tests were performed. * P<0.05.

### Confocal imaging experiments and analysis

For imaging experiments, U2OS PINK1 KO cells seeded at a density of 40,000 cells per 18mm glass coverslip in a 12 well plate were transiently transfected with 0.5 ug of plasmid DNA expressing WT-HA PINK1 under the control of a CMV weakened promoter using FuGene HD (Promega, E2311) at a ratio of 1:3. The media was changed after 2 h of transfection, and 20uM CCCP treatment was performed 48 h post-transfection for a period of 3 h in the absence of serum. The coverslips were fixed and stained for endogenous Tom20 (Santa Cruz Biotechnology, sc-17764) and exogenous HA (Biolegend, 902301), detected with donkey anti-mouse Alexa 488 (Thermo Fisher Scientific, A21202) and donkey anti-rabbit Alexa 555 (Thermo Fisher Scientific, A31572) secondary antibodies. Nuclei were counterstained with Hoechst (Thermo Fisher Scientific, H3570). Fluorescence images were processed using FIJI (ImageJ, NIH). The Mander’s Coefficients were obtained using JACoP. The mean difference in the Mander’s Coefficient between CCCP-treated and untreated cells was plotted in R. Changes relative to the WT were compared via general linear hypotheses tests as previously (79).

### Immunoprecipitation experiments

For FLAG immunopurification experiments, mock transfected HEK293T cells and PINK1-3xFLAG expressing cells were treated with 20 µM CCCP for 4 h followed by mitochondrial isolation and immunocapture using Anti-FLAG M2 Affinity gel. Bound proteins were eluted with FLAG peptides-containing elution buffer (containing 0.2% digitonin) and various fractions as indicated were subjected to BN-PAGE or SDS-PAGE followed by immunoblotting using PINK1, Tom22, Tom40 and FLAG antibodies.

For HA-immunopurification experiments, mock transfected U2OS PINK1 KO and PINK1-HA transfected cells were treated with 20 µM CCCP for 4 h followed by mitochondrial isolation and immunocapture using Pierce™ Anti-HA Magnetic Beads. Bound proteins were eluted with Pierce HA peptides (final concentration 2mg/ml)-containing elution buffer (containing 0.2% digitonin) and various fractions as indicated were subjected to BN-PAGE or SDS-PAGE followed by immunoblotting using PINK1, Tom22, Tom40, and Bak antibodies.

For His-immunopurification experiments, mock transfected HEK293T cells and PINK1-HIS transfected HEK293T cells were treated with 20 µM CCCP for 4 h followed by mitochondrial isolation and immunocapture using His CoPur affinity resin. Bound proteins were eluted with HIS-Select® Elution Buffer (containing 0.2% digitonin) and various fractions as indicated were subjected to BN-PAGE or SDS-PAGE followed by immunoblotting using PINK1, Tom22, and Tom40 antibodies.

### HMW PINK1 complex purification for mass spectrometry

HEK293T cells were mock transfected or transfected in triplicate with 30 µg of pTT5-PINK1-2xStrep-His (denoted as PINK1-Strep) using Lipofectamine 3000 according to manufacturer’s instructions. After 36 h, cells were treated with 20 µM CCCP for 3 h, and then harvested and subjected to mitochondrial isolation by nitrogen cavitation (as described above). Mitochondrial pellets were solubilized in lysis buffer with 1 % digitonin (25 mM HEPES pH 7.5, 150 mM NaCl, 5 % glycerol, 1 mM TCEP, supplemented with PhosStop (Roche), cOmplete protease inhibitor cocktail EDTA-free (Roche), and 1 mM EDTA). Lysates were incubated for 90 mins with end-over-end mixing and were spun at 15,000 rpm for 20 mins. Supernatants were diluted to 0.5% digitonin and were loaded onto Strep-Tactin resin (IBA). The resin was washed 3x with 25 mM HEPES pH 7.5, 150 mM NaCl, 5 % glycerol, 1 mM TCEP, 1 mM EDTA and 0.1 % digitonin. Samples were eluted in wash buffer containing 10 mM desthiobiotin. Eluted fractions were pooled, and proteins were precipitated using methanol-chloroform extraction. Pellets were re-solubilized in 8 M urea + 0.1 % ProteaseMAX surfactant (Promega) and were reduced with DTT and alkylated with iodoacetamide according to manufacturer’s instructions. 200 ng of Trypsin/Lys-C mix (Promega) was added, and the digestion was incubated at 37°C overnight. Trifluoroacetic acid was added to a final concentration of 0.5% to stop the digestion and samples were dried in a LabConco Centrivap DNA vacuum concentrator. Dried pellets were re-suspended in 0.5% formic acid / 5 % acetonitrile and were cleaned using C18 spin columns (Pierce), according to manufacturer’s instructions. Eluates were again dried in a LabConco Centrivap DNA vacuum concentrator.

### Mass spectrometry analysis

Extracted peptides were re-solubilized in 0.1% aqueous formic acid / 2% acetonitrile and loaded onto a Thermo Acclaim Pepmap (Thermo, 75 μm ID X 2 cm C18 3 μm beads) pre-column and then onto an Acclaim Pepmap EASY-Spray (Thermo, 75 μm X 15 cm with 2 μm C18 beads) analytical column separation using a Dionex Ultimate 3000 uHPLC at 250 nl/min with a gradient of 2–35% organic (0.1% formic acid in acetonitrile) over 1 hour. Peptides were analyzed using a Thermo Orbitrap Fusion mass spectrometer operating at 120,000 resolutions (FWHM in MS1) with HCD sequencing (15,000 resolution) at top speed for all peptides with a charge of 2 + or greater.

The raw data were processed using Andromeda, integrated into MaxQuant (version 2.1.4.0). Searches for tryptic (before K, R) peptides were performed against the H. sapiens proteome, with a minimum peptide length of 7 a.a. and a maximum of two missed cleavages. Cysteine carbamidomethylation was selected as a fixed modification. Protein N-term acetylation, methionine oxidation, phosphorylation (STY), and deamidation (NQ) were chosen as variable modifications. Default Orbitrap instrument parameters were used, including a first search peptide mass tolerance of 20 ppm and main search peptide tolerance of 4.5 ppm. Peptide spectral match and protein false discovery rate thresholds were set to 1 %. Label-free quantification (LFQ) was performed using only unique peptides for quantification. Match Between Runs was also used with default values.

MaxQuant LFQ values were analyzed using LFQ-Analyst as previously described (80). To summarize, data was pre-filtered to remove contaminants, reverse sequences, proteins only “identified by site”, proteins quantified with only a single peptide, and proteins with a high proportion of missing values. All LFQ values were transformed to a log2 scale, and missing values were imputed using the “Missing Not at Random” (MNAR) method, frequently used in Perseus software (81). Protein-wise linear models combined with empirical Bayes statistics were used for the differential expression analyses. Volcano plots were generated in GraphPad Prism using p-values and fold enrichments from LFQ-Analyst and iBAQ values from MaxQuant.

### AlphaFold multimer predictions on mass spectrometry hits

Iterative AlphaFold predictions for PINK1 against significantly enriched mass spectrometry hits were run on a local implementation of ColabFold Multimer using the AlphaFold framework (82, 83). Briefly, the full length PINK1 sequence was run against the sequences of the top 30 most enriched proteins with p-values < 0.05. Multiple sequence alignments were generated using mmseqs and used as input for the structure search using “colabfold_batch”. Parameters were set to 20 recycles with an RMSD tolerance of 0.5 Å, as previously described (84). PINK1:prey protein complex predictions were ranked by their average multimer score across the top 3 generated models and were visualized as a heat map in GraphPad Prism.

### Tim50 CRISPRi cloning and transfection

The dCas9 plasmid used was pLX_311-KRAB-dCas9 (Addgene #96918, gift from John Doench & William Hahn & David Root; referred to as CRISPRi). gRNA plasmid cloning was adapted from Weissman lab protocols, using the pCRISPRi/a-v2 plasmid (Addgene #84832, gift from Jonathan Weissman). Two sgRNA’s were used to target TIMM50, sgRNA1 (5’-GTCCGGGACGCCTCACCTCA-3’), and sgRNA2 (5’-GTGGCGTCAGCGCAAGATGG-3’). For transfection of CRISPRi machinery and sgRNA plasmids, ∼5×10^6^ HEK293T cells were seeded in a 150 mm dish, and after 24 h were co-transfected with dCas9, sgRNA1 and sgRNA2 in equimolar amounts using jetPRIME transfection reagent (Polyplus). Transfection was performed according to manufacturer’s instructions, and cells were harvested 36 h post-transfection for following mitochondrial isolation and SDS-PAGE and BN-PAGE immunoblotting.

## Author contributions

MAE, ANB, JFT & EAF conceived and designed the study. MAE, ANB, AF, EJM, TG, AS, CEZ, JFT performed research, analyzed and interpreted the data. MAE, AF, TG, RAT, AS, NK and TMD provided new reagents. MAE, ANB, JFT and EAF wrote the manuscript with input from all authors.

## Acknowledgments

MAE is CIHR Banting Fellow and was supported by Parkinson Canada. TG was supported by Parkinson Canada. ANB was supported by a CIHR Doctoral Fellowship and a CRBS Maximilian Eivaskhani In Memoriam Graduate Studentship. We thank Amy Wong and Lorne Taylor at the Proteomics and Molecular Analysis Platform for LC-MS data acquisition.

---

**Supplemental Figure 1.**
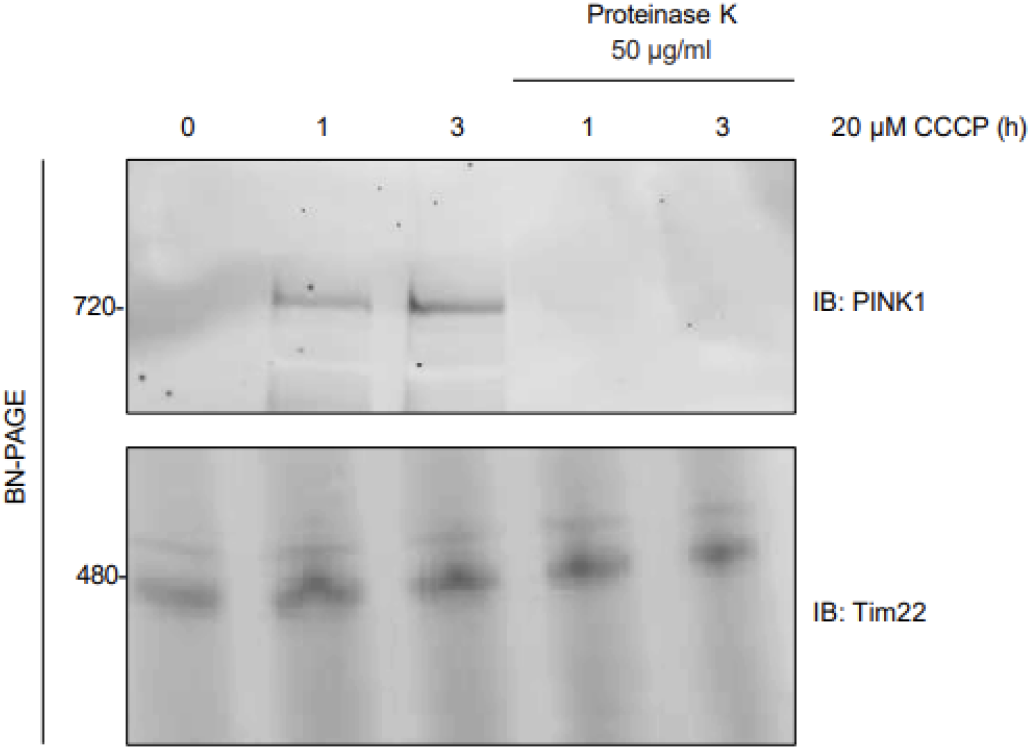
Mitochondria were isolated from U2OS WT cells that were either treated with CCCP or vehicle control (DMSO). Isolated mitochondria were treated with or without external protease (PK, 20mg/ml) solubilized in 1% digitonin containing buffer and subjected to BN-PAGE and immunoblotting using antibodies against PINK1 (outer membrane), and TIM22 complex (inner membrane).

**Supplemental Figure 2.**
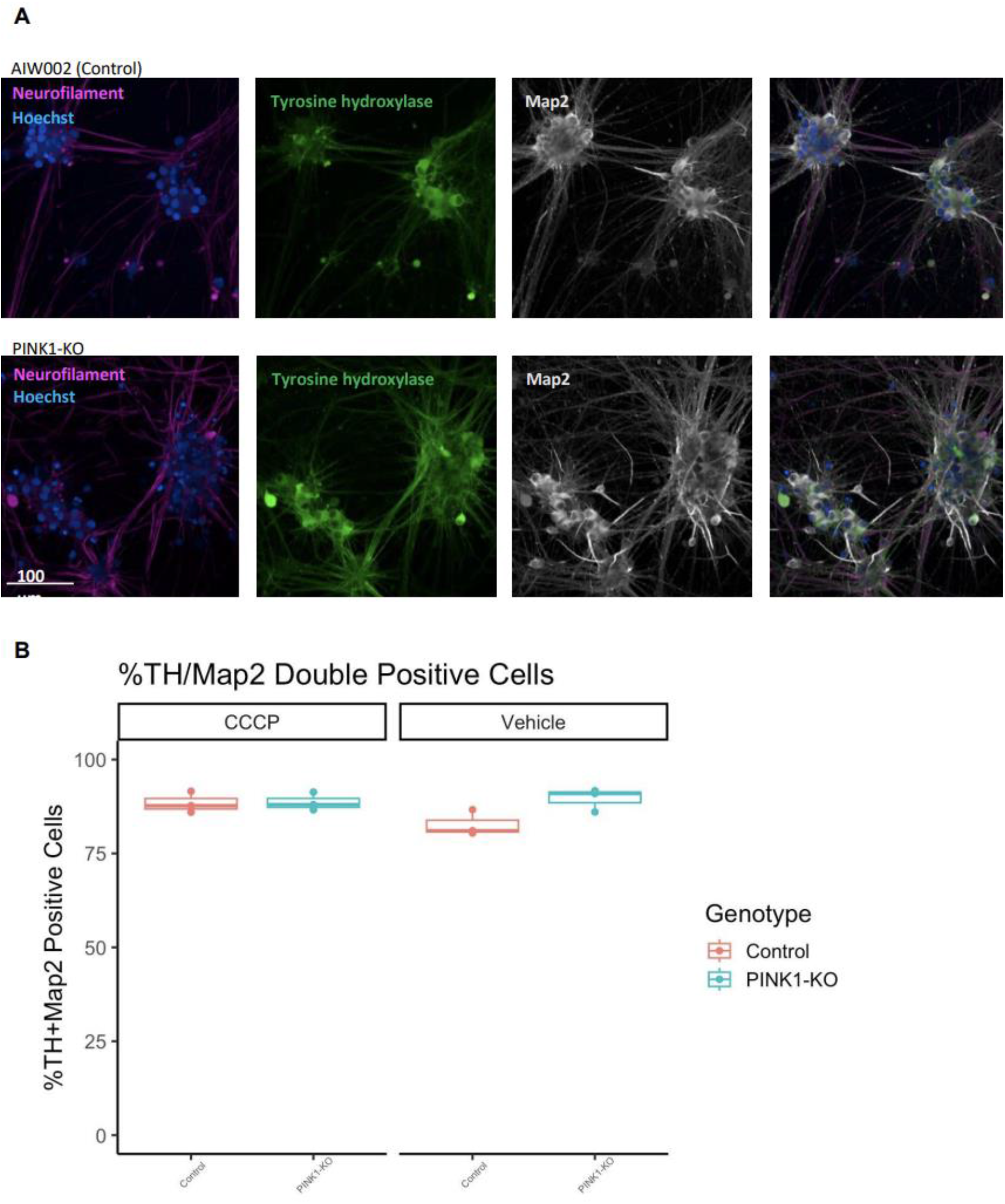
Neuronal identity was assessed by immunofluorescent staining for Map2 (neurons) and tyrosine hydroxylase (TH - dopaminergic neurons). After 4 weeks of differentiation neurons were fixed with 4% paraformaldehyde for 20 mins, permeabilized for 10 mins with 0.3% saponin in PBS, and blocked for 1 h with 1% BSA and 4% goat-serum in PBS. Primary antibodies against TH (Pelfreeze, P40101) and Map2 (Encor, CPCA-MAP2) were diluted (1:500 and 1:2000 respectively) in blocking solution and incubated at 4 degrees overnight. Neurons were then washed 3 times in PBS and incubated for 1 hour with AlexaFluor secondary antibodies in blocking solution. Neurons were then washed twice and stained with Hoechst before imaging. Imaging was performed on an Opera Phenix high content confocal microscope using 20X water immersion objective. Image analysis was performed using the Columbus software and data processing was then conducted using R studio.

**Supplemental Figure 3.**
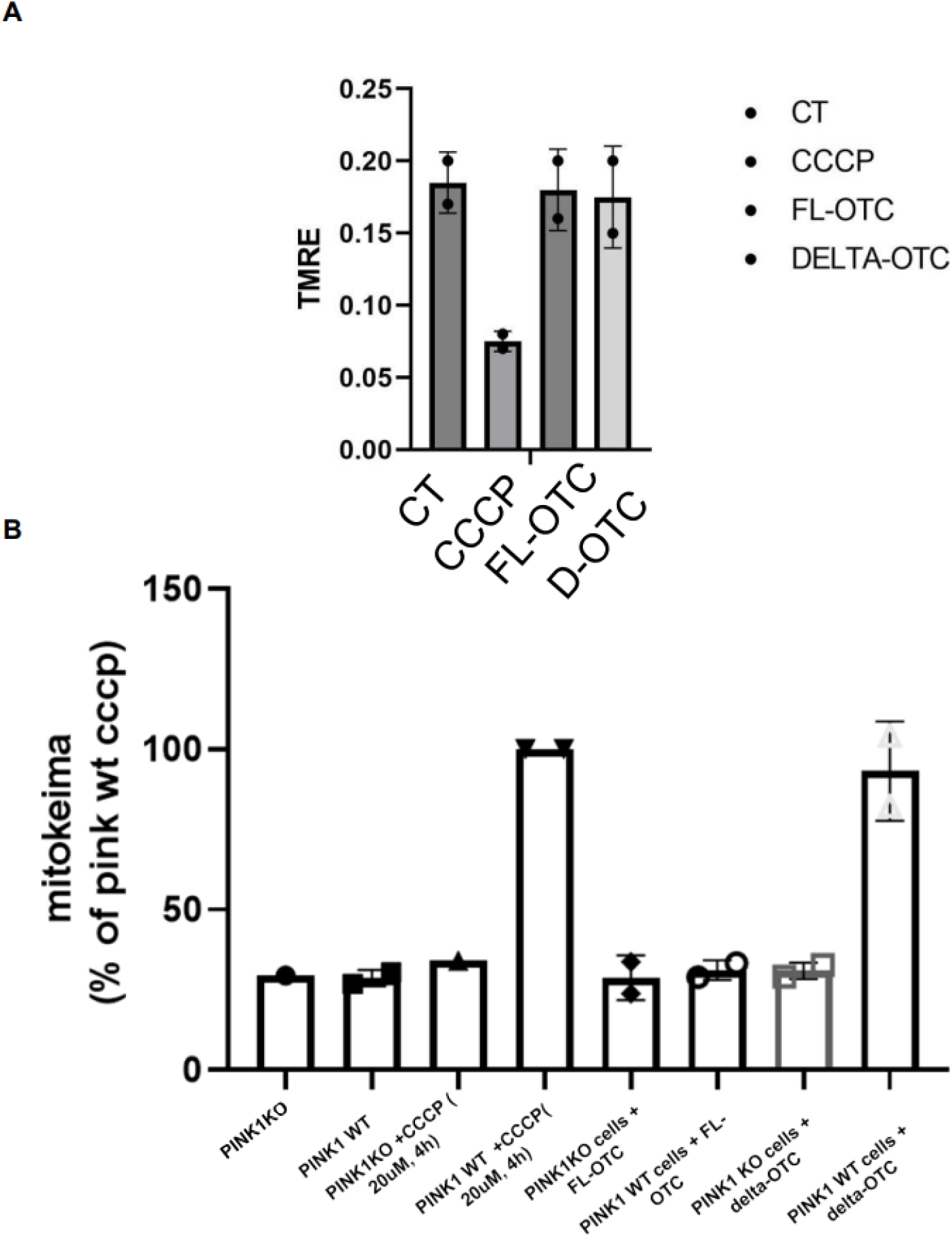
A) ΔOTC, full length OTC (FL-OTC) expressing cells and cells treated with 20 µM CCCP (4 h) cells were detached, stained with TMRE, and analyzed for the intensity of TMRE using a fluorescence plate reader as recommended by the manufacturer. As a negative control, untreated cells were stained with TMRE before FACS analysis. FACS results were represented as mean ± SEM from three independent experiments. B) U2OS cells transfected with pCMV(d1) PINK1 (WT or the indicated constructs as FL-OTC or ΔOTC) were treated with 20 μM CCCP or DMSO for 4 h and were subjected to mt-Keima reporter assays. Bars indicate the relative level of mitophagy, normalized to WT PINK1 treated with CCCP, plotted as mean (n = 2).

**Supplemental Figure 4.**
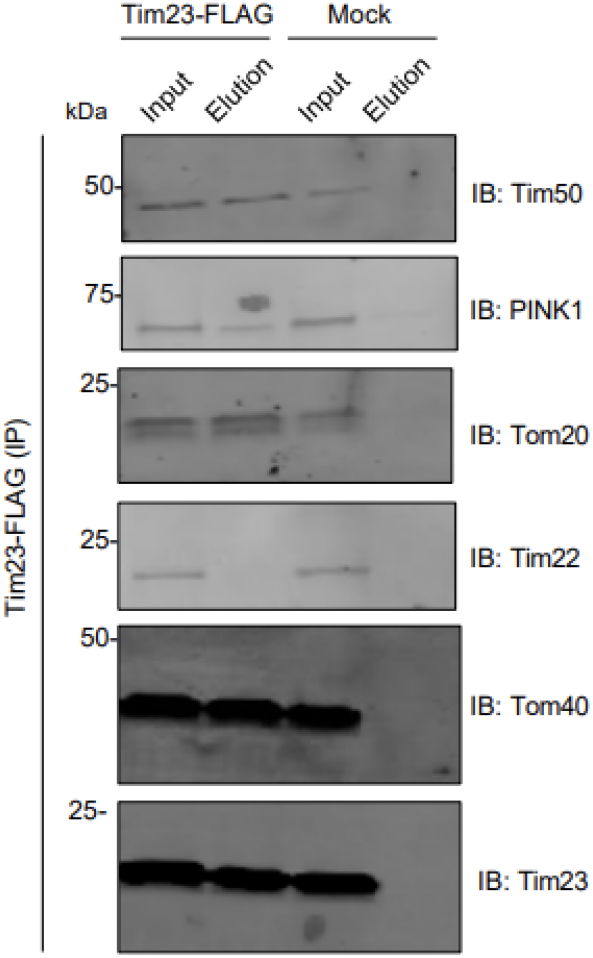
Mock or Tim23-FLAG transfected HEK293T cells were treated with 20 µM CCCP for 4 h followed by mitochondrial isolation and immunocapture using M2 FLAG affinity gel. Bound proteins were eluted with FLAG peptide and various fractions were subjected to SDS-PAGE immunoblotting using the indicated antibodies.

**Supplemental Figure 5.**
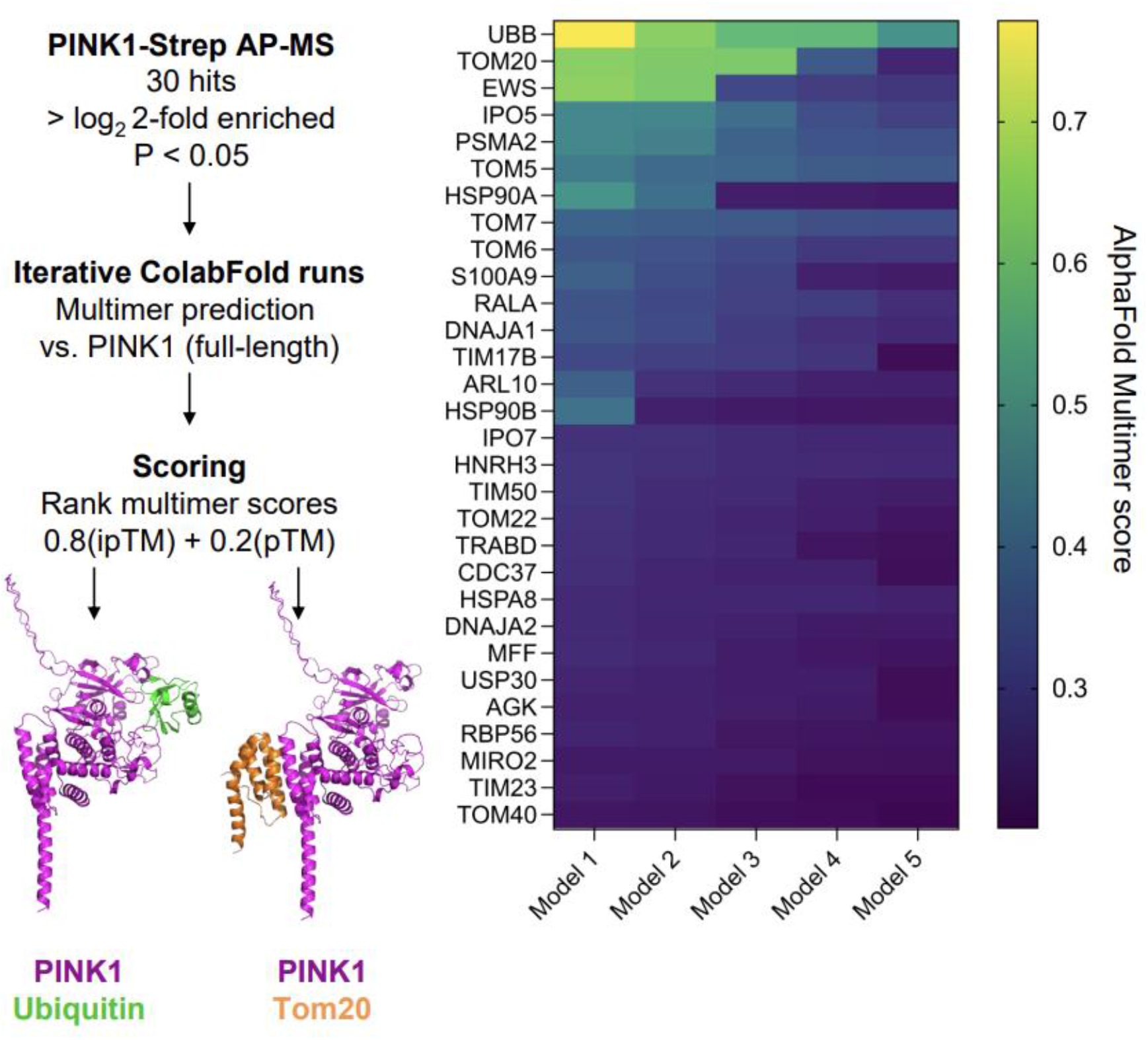
ColabFold multimer was run locally using full-length PINK1 as a bait against all enriched mass spectrometry hits. Complexes were sorted according to the average of their Multimer scores across the top 3 models. PINK1:Tom20 and PINK1:Ub structures were visualized in PyMOL.

**Supplemental Figure 6.**
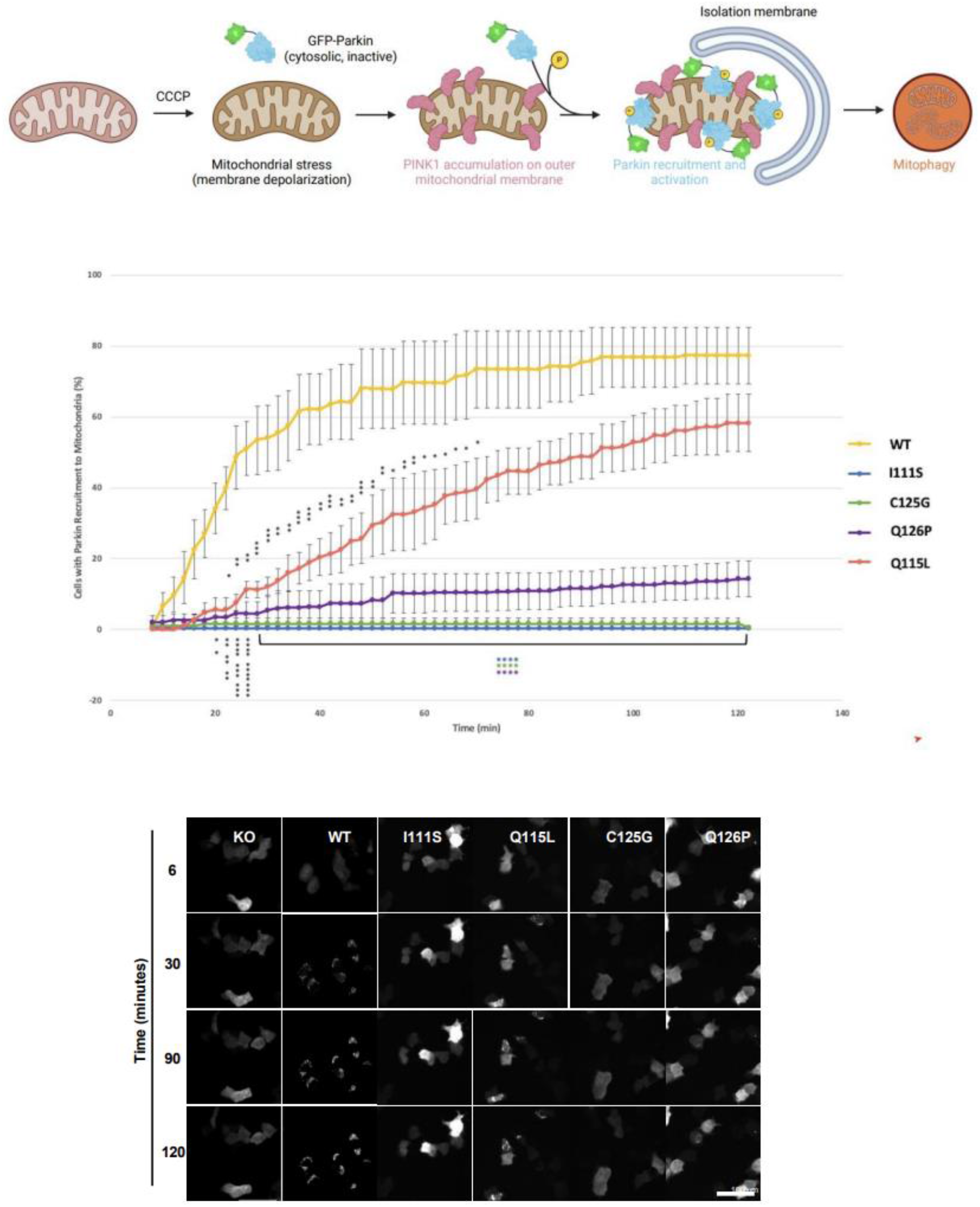
Time-lapse imaging of parkin recruitment to mitochondria upon treatment with 20 µM CCCP in U2OS PINK1 KO cells co-expressing with WT eGFP-Parkin and α-NTE PINK1 mutants (I111S, Q115L, C125G, Q126P) as indicated. Recruitment can be visualized by the appearance of punctate GFP fluorescence. Scale bar: 100 µm.

**Supplemental Figure 7.**
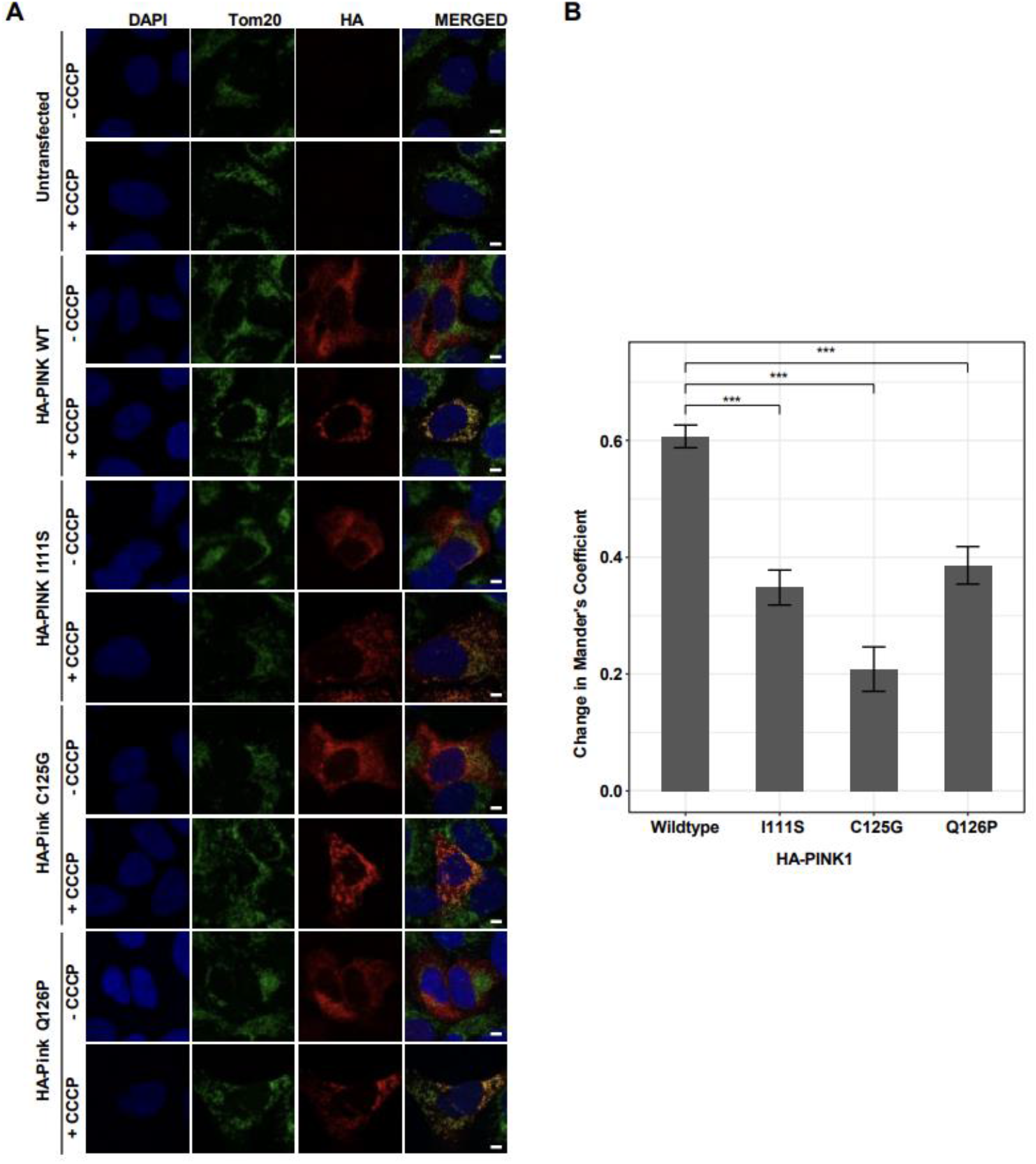
Confocal microscopy of PINK KO U2OS cells that were either untransfected or transiently transfected with the indicated PINK1-HA constructs with or without CCCP treatment. The scale bar represents 5 μm. B. The bars indicate the mean differences in the Mander’s Coefficient between CCCP-treated and untreated cells, and the standard error of the mean differences. Changes relative to the WT were compared via general linear hypotheses tests (*** all adjusted *p* < 1e-7; *t* > 5.82).

**Supplemental Figure 8.**
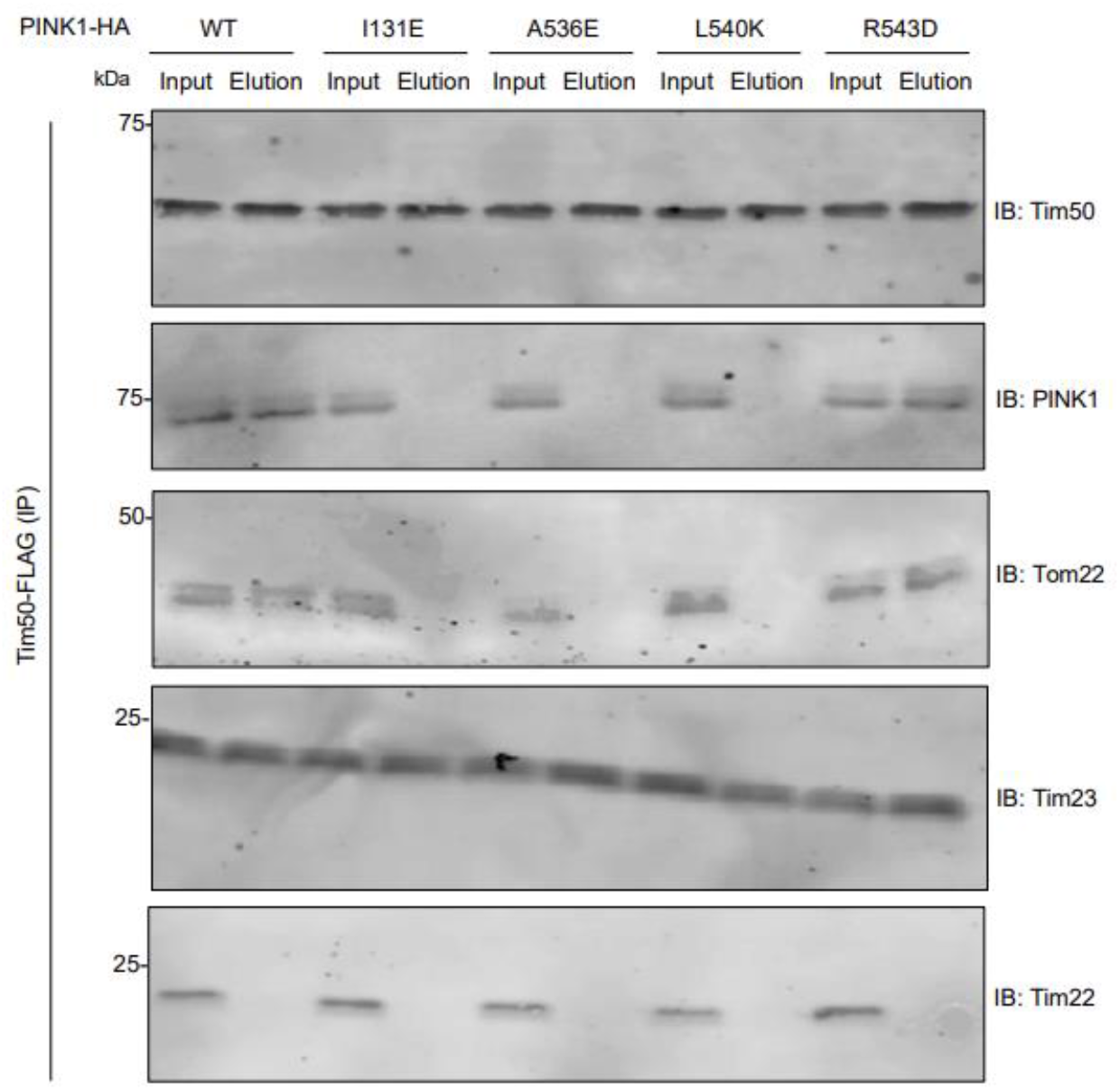
U2OS PINK1 KO cells were co-transfected with indicated PINK1 mutants and Tim50-FLAG for 48 h, treated with 20 µM CCCP for 4 h and mitochondria were isolated by N_2_ cavitation. Mitochondrial pellets were lysed in 1 % digitonin, and lysates were subjected to immunocapture using Anti-FLAG M2 Affinity gel. Bound proteins were eluted with FLAG peptides-containing elution buffer (containing 0.2% digitonin) and various fractions as indicated were subjected to SDS-PAGE followed by immunoblotting using the indicated antibodies.

## References

1. S. G. Reich, J. M. Savitt, Parkinson’s Disease. Med. Clin. North Am. 103, 337–350 (2019).

2. A. M. Pickrell, R. J. Youle, The roles of PINK1, parkin, and mitochondrial fidelity in Parkinson’s disease. Neuron 85, 257–273 (2015).

3. I. Martin, V. L. Dawson, T. M. Dawson, Recent Advances in the Genetics of Parkinson’s Disease. Annu. Rev. Genomics Hum. Genet. 12, 301–325 (2011).

4. M. A. Eldeeb, R. A. Thomas, M. A. Ragheb, A. Fallahi, E. A. Fon, Mitochondrial quality control in health and in Parkinson’s disease. Physiol. Rev. 102, 1721–1755 (2022).

5. S. Pickles, P. Vigié, R. J. Youle, Mitophagy and Quality Control Mechanisms in Mitochondrial Maintenance. Curr. Biol. CB 28, R170–R185 (2018).

6. A. B. Harbauer, R. P. Zahedi, A. Sickmann, N. Pfanner, C. Meisinger, The protein import machinery of mitochondria-a regulatory hub in metabolism, stress, and disease. Cell Metab. 19, 357–372 (2014).

7. L. Wrobel, et al., Mistargeted mitochondrial proteins activate a proteostatic response in the cytosol. Nature 524, 485–488 (2015).

8. X. Wang, X. J. Chen, A cytosolic network suppressing mitochondria-mediated proteostatic stress and cell death. Nature 524, 481–484 (2015).

9. F. Boos, et al., Mitochondrial protein-induced stress triggers a global adaptive transcriptional programme. Nat. Cell Biol. 21, 442–451 (2019).

10. H. Weidberg, A. Amon, MitoCPR-A surveillance pathway that protects mitochondria in response to protein import stress. Science 360, eaan4146 (2018).

11. C. U. Mårtensson, et al., Mitochondrial protein translocation-associated degradation. Nature 569, 679–683 (2019).

12. E. M. Valente, et al., Hereditary early-onset Parkinson’s disease caused by mutations in PINK1. Science 304, 1158–1160 (2004).

13. T. Kitada, et al., Mutations in the parkin gene cause autosomal recessive juvenile parkinsonism. Nature 392, 605–608 (1998).

14. N. Matsuda, et al., PINK1 stabilized by mitochondrial depolarization recruits Parkin to damaged mitochondria and activates latent Parkin for mitophagy. J. Cell Biol. 189, 211–221 (2010).

15. K. Shiba-Fukushima, et al., Phosphorylation of Mitochondrial Polyubiquitin by PINK1 Promotes Parkin Mitochondrial Tethering. PLoS Genet. 10, e1004861 (2014).

16. D. Narendra, A. Tanaka, D.-F. Suen, R. J. Youle, Parkin is recruited selectively to impaired mitochondria and promotes their autophagy. J. Cell Biol. 183, 795–803 (2008).

17. F. Koyano, et al., Ubiquitin is phosphorylated by PINK1 to activate parkin. Nature 510, 162–166 (2014).

18. M. Lazarou, S. M. Jin, L. A. Kane, R. J. Youle, Role of PINK1 binding to the TOM complex and alternate intracellular membranes in recruitment and activation of the E3 ligase Parkin. Dev. Cell 22, 320–333 (2012).

19. K. Okatsu, et al., A dimeric PINK1-containing complex on depolarized mitochondria stimulates Parkin recruitment. J. Biol. Chem. 288, 36372–36384 (2013).

20. L. A. Kane, et al., PINK1 phosphorylates ubiquitin to activate Parkin E3 ubiquitin ligase activity. J. Cell Biol. 205, 143–153 (2014).

21. S. Sekine, R. J. Youle, PINK1 import regulation; a fine system to convey mitochondrial stress to the cytosol. BMC Biol. 16, 2 (2018).

22. A. N. Bayne, J.-F. Trempe, Mechanisms of PINK1, ubiquitin and Parkin interactions in mitochondrial quality control and beyond. Cell. Mol. Life Sci. CMLS 76, 4589–4611 (2019).

23. A. Tanaka, et al., Proteasome and p97 mediate mitophagy and degradation of mitofusins induced by Parkin. J. Cell Biol. 191, 1367–1380 (2010).

24. S. Geisler, et al., PINK1/Parkin-mediated mitophagy is dependent on VDAC1 and p62/SQSTM1. Nat. Cell Biol. 12, 119–131 (2010).

25. N. C. Chan, et al., Broad activation of the ubiquitin-proteasome system by Parkin is critical for mitophagy. Hum. Mol. Genet. 20, 1726–1737 (2011).

26. L. Glauser, S. Sonnay, K. Stafa, D. J. Moore, Parkin promotes the ubiquitination and degradation of the mitochondrial fusion factor mitofusin 1. J. Neurochem. 118, 636–645 (2011).

27. A. Sugiura, G.-L. McLelland, E. A. Fon, H. M. McBride, A new pathway for mitochondrial quality control: mitochondrial-derived vesicles. EMBO J. 33, 2142–2156 (2014).

28. A. F. Schubert, et al., Structure of PINK1 in complex with its substrate ubiquitin. Nature 552, 51–56 (2017).

29. A. B. Harbauer, et al., Neuronal mitochondria transport Pink1 mRNA via synaptojanin 2 to support local mitophagy. Neuron 110, 1516–1531.e9 (2022).

30. A. W. Greene, et al., Mitochondrial processing peptidase regulates PINK1 processing, import and Parkin recruitment. EMBO Rep. 13, 378–385 (2012).

31. S. M. Jin, et al., Mitochondrial membrane potential regulates PINK1 import and proteolytic destabilization by PARL. J. Cell Biol. 191, 933–942 (2010).

32. S. Sekine, et al., Reciprocal Roles of Tom7 and OMA1 during Mitochondrial Import and Activation of PINK1. Mol. Cell 73, 1028–1043.e5 (2019).

33. K. Yamano, R. J. Youle, PINK1 is degraded through the N-end rule pathway. Autophagy 9, 1758– 1769 (2013).

34. M. Ando, et al., The PINK1 p.I368N mutation affects protein stability and ubiquitin kinase activity. Mol. Neurodegener. 12, 32 (2017).

35. D. P. Narendra, et al., PINK1 Is Selectively Stabilized on Impaired Mitochondria to Activate Parkin. PLOS Biol. 8, e1000298 (2010).

36. S. A. Hasson, et al., High-content genome-wide RNAi screens identify regulators of parkin upstream of mitophagy. Nature 504, 291–295 (2013).

37. S. Rasool, et al., Mechanism of PINK1 activation by autophosphorylation and insights into assembly on the TOM complex. Mol. Cell 82, 44–59.e6 (2022).

38. P. Kakade, et al., Mapping of a N-terminal α-helix domain required for human PINK1 stabilization, Serine228 autophosphorylation and activation in cells. Open Biol. 12, 210264 (2022).

39. S. Akabane, et al., TIM23 facilitates PINK1 activation by safeguarding against OMA1-mediated degradation in damaged mitochondria. Cell Rep. 42, 112454 (2023).

40. L. Bolliger, T. Junne, G. Schatz, T. Lithgow, Acidic receptor domains on both sides of the outer membrane mediate translocation of precursor proteins into yeast mitochondria. EMBO J. 14, 6318–6326 (1995).

41. M. Moczko, et al., The intermembrane space domain of mitochondrial Tom22 functions as a trans binding site for preproteins with N-terminal targeting sequences. Mol. Cell. Biol. 17, 6574 (1997).

42. D. Rapaport, W. Neupert, R. Lill, Mitochondrial protein import. Tom40 plays a major role in targeting and translocation of preproteins by forming a specific binding site for the presequence. J. Biol. Chem. 272, 18725–18731 (1997).

43. T. Komiya, et al., Interaction of mitochondrial targeting signals with acidic receptor domains along the protein import pathway: evidence for the ‘acid chain’ hypothesis. EMBO J. 17, 3886–3898 (1998).

44. T. Kanamori, et al., Uncoupling of transfer of the presequence and unfolding of the mature domain in precursor translocation across the mitochondrial outer membrane. Proc. Natl. Acad. Sci. 96, 3634–3639 (1999).

45. H. Schägger, G. von Jagow, Blue native electrophoresis for isolation of membrane protein complexes in enzymatically active form. Anal. Biochem. 199, 223–231 (1991).

46. N.-V. Mohamed, et al., Midbrain organoids with an SNCA gene triplication model key features of synucleinopathy. Brain Commun. 3, fcab223 (2021).

47. C. X.-Q. Chen, et al., Generation of homozygous PRKN, PINK1 and double PINK1/PRKN knockout cell lines from healthy induced pluripotent stem cells using CRISPR/Cas9 editing. Stem Cell Res. 62, 102806 (2022).

48. S. M. Jin, R. J. Youle, The accumulation of misfolded proteins in the mitochondrial matrix is sensed by PINK1 to induce PARK2/Parkin-mediated mitophagy of polarized mitochondria. Autophagy 9, 1750–1757 (2013).

49. Q. Zhao, et al., A mitochondrial specific stress response in mammalian cells. EMBO J. 21, 4411– 4419 (2002).

50. M. U. Stevens, et al., Structure-based design and characterization of Parkin activating mutations. 2022.02.22.481412 (2022).

51. H. Kato, Q. Lu, D. Rapaport, V. Kozjak-Pavlovic, Tom70 Is Essential for PINK1 Import into Mitochondria. PLOS ONE 8, e58435 (2013).

52. K. K. Maruszczak, M. Jung, S. Rasool, J.-F. Trempe, D. Rapaport, The role of the individual TOM subunits in the association of PINK1 with depolarized mitochondria. J. Mol. Med. 100, 747–762 (2022).

53. M. Bhagawati, et al., The receptor subunit Tom20 is dynamically associated with the TOM complex in mitochondria of human cells. Mol. Biol. Cell 32, br1 (2021).

54. J. Su, et al., Structural basis of Tom20 and Tom22 cytosolic domains as the human TOM complex receptors. Proc. Natl. Acad. Sci. 119, e2200158119 (2022).

55. W. Wang, et al., Atomic structure of human TOM core complex. Cell Discov. 6, 1–10 (2020).

56. E. Deas, H. Plun-Favreau, N. W. Wood, PINK1 function in health and disease. EMBO Mol. Med. 1, 152–165 (2009).

57. R. Marongiu, et al., Whole gene deletion and splicing mutations expand the PINK1 genotypic spectrum. Hum. Mutat. 28, 98 (2007).

58. P. Ibáñez, et al., Mutational analysis of the PINK1 gene in early-onset parkinsonism in Europe and North Africa. Brain 129, 686–694 (2006).

59. J. Prestel, et al., Clinical and molecular characterisation of a Parkinson family with a novel PINK1 mutation. J. Neurol. 255, 643–648 (2008).

60. A. M. Schlitter, et al., Exclusion of PINK1 as candidate gene for the late-onset form of Parkinson’s disease in two European populations. J. Negat. Results Biomed. 4, 10 (2005).

61. M. J. Landrum, et al., ClinVar: improving access to variant interpretations and supporting evidence. Nucleic Acids Res. 46, D1062–D1067 (2018).

62. C. Zhang, et al., The plant triterpenoid celastrol blocks PINK1-dependent mitophagy by disrupting PINK1’s association with the mitochondrial protein TOM20. J. Biol. Chem. 294, 7472–7487 (2019).

63. A. Chacinska, et al., Mitochondrial Presequence Translocase: Switching between TOM Tethering and Motor Recruitment Involves Tim21 and Tim17. Cell 120, 817–829 (2005).

64. R. Gomkale, et al., Mapping protein interactions in the active TOM-TIM23 supercomplex. Nat. Commun. 12, 5715 (2021).

65. S. I. Sim, Y. Chen, D. L. Lynch, J. C. Gumbart, E. Park, Structural basis of mitochondrial protein import by the TIM23 complex. Nature, 1–7 (2023).

66. X. Zhou, et al., Molecular pathway of mitochondrial preprotein import through the TOM-TIM23 supercomplex. 2023.06.21.546012 (2023).

67. V. A. M. Gold, et al., Visualizing active membrane protein complexes by electron cryotomography. Nat. Commun. 5, 4129 (2014).

68. K. Tucker, E. Park, Cryo-EM structure of the mitochondrial protein-import channel TOM complex at near-atomic resolution. Nat. Struct. Mol. Biol. 26, 1158–1166 (2019).

69. K. Okatsu, et al., PINK1 autophosphorylation upon membrane potential dissipation is essential for Parkin recruitment to damaged mitochondria. Nat. Commun. 3, 1016 (2012).

70. S. Rasool, et al., PINK1 autophosphorylation is required for ubiquitin recognition. EMBO Rep. 19, e44981 (2018).

71. M. Y. Tang, et al., Structure-guided mutagenesis reveals a hierarchical mechanism of Parkin activation. Nat. Commun. 8, 14697 (2017).

72. C. X.-Q. Chen, et al., A Multistep Workflow to Evaluate Newly Generated iPSCs and Their Ability to Generate Different Cell Types. Methods Protoc. 4, 50 (2021).

73. N.-V. Mohamed, et al., Microfabricated disk technology: Rapid scale up in midbrain organoid generation. Methods San Diego Calif 203, 465–477 (2022).

74. X. Chen, C. Rocha, T. Rao, T. M. Durcan, NeuroEDDU protocols_iPSC culture (2019) https://doi.org/10.5281/zenodo.3738269 (July 14, 2023).

75. X. Chen, et al., Induction of Dopaminergic or Cortical neuronal progenitors from iPSCs (2019) https://doi.org/10.5281/zenodo.3738358 (July 14, 2023).

76. X. Chen, N. Lauinger, C. Rocha, T. Rao, T. M. Durcan, Generation of dopaminergic or cortical neurons from neuronal progenitors (2019) https://doi.org/10.5281/zenodo.3738323 (July 14, 2023).

77. W. Yi, et al., The landscape of Parkin variants reveals pathogenic mechanisms and therapeutic targets in Parkinson’s disease. Hum. Mol. Genet. 28, 2811–2825 (2019).

78. V. M. Fava, et al., Pleiotropic effects for Parkin and LRRK2 in leprosy type-1 reactions and Parkinson’s disease. Proc. Natl. Acad. Sci. U. S. A. 116, 15616–15624 (2019).

79. T. Hothorn, F. Bretz, P. Westfall, Simultaneous inference in general parametric models. Biom. J. Biom. Z. 50, 346–363 (2008).

80. A. D. Shah, R. J. A. Goode, C. Huang, D. R. Powell, R. B. Schittenhelm, LFQ-Analyst: An Easy-To-Use Interactive Web Platform To Analyze and Visualize Label-Free Proteomics Data Preprocessed with MaxQuant. J. Proteome Res. 19, 204–211 (2020).

81. S. Tyanova, T. Temu, J. Cox, The MaxQuant computational platform for mass spectrometry-based shotgun proteomics. Nat. Protoc. 11, 2301–2319 (2016).

82. M. Mirdita, et al., ColabFold: making protein folding accessible to all. Nat. Methods 19, 679–682 (2022).

83. J. Jumper, et al., Highly accurate protein structure prediction with AlphaFold. Nature 596, 583–589 (2021).

84. R. Yin, B. Y. Feng, A. Varshney, B. G. Pierce, Benchmarking AlphaFold for protein complex modeling reveals accuracy determinants. Protein Sci. 31, e4379 (2022).

